# Biomarkers to distinguish bacterial from viral pediatric clinical pneumonia in a malaria endemic setting

**DOI:** 10.1101/2020.04.27.036277

**Authors:** Michael A. Gillette, D. R. Mani, Christopher Uschnig, Karell G. Pellé, Lola Madrid, Sozinho Acácio, Miguel Lanaspa, Pedro Alonso, Clarissa Valim, Steven A. Carr, Stephen F. Schaffner, Bronwyn MacInnis, Danny A. Milner, Quique Bassat, Dyann F. Wirth

**Author notes:** To whom correspondence should be addressed: Dyann F. Wirth. Co-Authorship, the co-authors contributed equally (Michael A. Gillette, D. R. Mani, Christopher Uschnig). Co-Authorship (Quique Bassat, Dyann F. Wirth).

## Abstract

**BACKGROUND:** Differentiating the etiology of acute febrile respiratory illness in children is a challenge in low-income, malaria-endemic settings because the main pathogens responsible (viruses, bacteria, and malaria parasites) overlap in clinical presentation and frequently occur together as mixed infections. The critical task is to rapidly identify bacterial pneumonia to enable appropriate antibiotic treatment, ideally at point of care. Current diagnostic tests are insufficient and there is a need for the discovery and development of new tools. Here we report the identification of a unique biomarker signature that can be identified in blood samples.

**METHODS:** Blood samples from 195 pediatric Mozambican patients with clinical pneumonia were analyzed with an aptamer-based high dynamic range assay to quantify ∼1200 proteins. For discovery of new biomarkers, we identified a training set of patient samples in which the underlying etiology of the pneumonia was established as bacterial, viral or malaria. Proteins whose abundances varied significantly between patients with verified etiologies (FDR<0.01) formed the basis for predictive diagnostic models that were created using machine learning techniques (Random Forest, Elastic Net). These models were validated on a dedicated test set of samples.

**RESULTS:** 219 proteins had significantly different abundances between bacterial and viral infections, and 151 differed between bacterial infections and a mixed pool of viral and malaria infections. Predictive diagnostic models achieved >90% sensitivity and >80% specificity, regardless of whether one or two pathogen classes were present. Bacterial pneumonia was strongly associated with markers of neutrophil activity, in particular neutrophil degranulation. Degranulation markers included HP, LCN2, LTF, MPO, MMP8, PGLYRP1, RETN, SERPINA1, S100A9, and SLPI.

**CONCLUSION:** Blood protein signatures highly associated with neutrophil biology reliably differentiated bacterial pneumonia from other causes. With appropriate technology, these markers could provide the basis for a rapid diagnostic for field-based triage for antibiotic treatment of pediatric pneumonia.

## Introduction

Febrile respiratory illness is a leading cause of mortality and morbidity among children around the world. Identifying the underlying cause — bacterial^1^, viral, or (less commonly) malaria^2,3^ — is crucially important but difficult, as clinical presentations of the three infections can be very similar. The critical need is to identify bacterial infections^3^, so they can be treated appropriately.^4,5^ Bacterial diagnosis is plagued by a lack of effective diagnostic tests: laborious microbiological culture or molecular testing methods, where available, are often not sensitive enough to detect the underlying bacterial pathogen^6^, and neither are radiological evaluations (through chest-X-ray or ultrasound) which seldom are accessible in low-resource settings. Heightening the diagnostic dilemma, it is becoming increasingly clear that malaria or viral infections and bacterial secondary co-infections occur commonly together.^7,8^

Cellular responses to bacterial, viral, and malaria infections are distinct, being chiefly neutrophilic, lymphocytic, or monocytic, respectively. Thus, host response signatures have been explored as possible diagnostic indicators. To date, however, these approaches have not proven to be sufficiently reliable^9–13^, based on a recommended benchmark^14^ of acceptable and desirable thresholds for sensitivity (desirable ≥95%, acceptable ≥90%) and specificity (≥90% and ≥80%) for diagnostic tests. We hypothesized that the distinctive cellular host responses could be detected at the protein level. Here we describe a test of this hypothesis, based on the differential expression of proteins in blood specimens collected from children with febrile respiratory illness in southern Mozambique, where malaria is endemic. Febrile respiratory illness cases were assigned by all available gold standard tests, and using highly specific case-definitions, to one of three underlying causes, bacteria, viruses, or malaria, or to a combination of those (“mixed infections”). Proteins were assayed with SOMAScan technology (Somalogic, Boulder, CO), an array-based modified aptamer platform with a broad representation of biological pathways including inflammation, signal transduction, and immune processes. This technology provides a quantitative assay of approximately 1200 proteins simultaneously, offers a high dynamic range, and has modest sample requirements (150 ul plasma).^15^

The resulting protein expression data were used to create machine learning-based models for distinguishing bacterial from viral or malaria infections. The same data, along with data from our prior RNA- and protein-based studies^9,13^, provided the basis for pathway analyses, to help confirm the underlying biology of the host response.

## Methods

### Study Design

The study recruited two groups of children under the age of 10 years at the Manhiça District Hospital in Mozambique. The first group included children with febrile respiratory illness admitted to the hospital fulfilling the “clinical pneumonia” criteria (as defined by WHO), and the second were afebrile and symptomless healthy community controls used to establish a baseline. Febrile respiratory illness cases were assigned by all available gold standard tests to one of three underlying causes, bacteria, viruses, or malaria, or to a combination of those (“mixed infections”).

### Study population and sample classification procedure

Children with documented fever at admission (>37.5°C axillary temperature) or a history of fever in the preceding 24 hours who met the WHO case definition for clinical pneumonia (increased respiratory rate and cough or difficulty breathing) ^16^were approached and recruited to the study. All children underwent anteroposterior chest radiography; X-ray images were independently interpreted following the WHO recommended guidelines for pneumonia diagnosis by two experienced clinicians.^17^ Informed consent was obtained from parents/guardians.

Patients were classified as having clinical pneumonia associated with bacterial (69 samples) malaria (42 samples) or viral (48 samples) infection using the criteria described in Valim et al. (2016), with minor modifications. In brief, patients were classified as bacterial pneumonia when pathogenic bacteria were isolated (or detected through RT-PCR) from blood or pleural exudate, and after confirming the absence of malarial infection. Viral pneumonia required the detection in the nasopharyngeal aspirate (NPA) of a viral respiratory pathogen, no isolated bacteria in the blood culture or RT-PCR, no “endpoint pneumonia” in the chest X-ray, and negative malaria microscopy. Finally, a malaria case required a positive malaria smear microscopy (according to pre-determined parasitemia thresholds in relation to age^18^), normal chest X-ray and no detectable bacterial infection.

To address the known insensitivity of blood culture for bacterial pneumonia, cases were also assigned a bacterial etiology if the NPA was negative for virus but the patient had leukocytosis and a dense radiographic consolidation (endpoint pneumonia) based on consensus of two independent experts. Since NPAs are often positive on RT-PCR for potential viral respiratory pathogens even in clinically well children, the detection of a virus in the nasopharyngeal aspirate did not alter the class assignments for confirmed bacterial or malarial cases. See Fig. S1 for a comprehensive flowchart for patient classification.

In addition, 23 patient samples with other mixed infections were also included in the study: Bacteria & Malaria (n=9), Virus & probable Bacterial secondary co-infection (n=11), Malaria & Virus (n=2), and Malaria & Other (n=1) (for details see Table S3). “Virus & probable bacterial secondary co-infection” samples were virus positive, culture and PCR-negative for bacteria but with leukocytosis and radiographic endpoint pneumonia, suggestive of a secondary bacterial infection.

### SOMAScan protein assay

The SOMAScan assay uses SOMAmers (Slow Off-rate Modified Aptamers) to capture proteins and translates binding events into signals measured in Relative Fluorescence Units (RFU). RFU are directly proportional to the abundance of the target proteins in the sample, as informed by a standard curve generated for each protein-SOMAmer pair. The dynamic range of the assay is enhanced by three serial dilutions of the sample, with the least concentrated dilution used to quantify the most abundant proteins (∼μM concentration in the original sample), and the most concentrated used for the least abundant proteins (fM to pM concentration).^15^ Samples were assayed in two batches; the SOMAScan assays used in the first set of 167 samples quantified 1129 proteins and the SOMAScan assay used in the second set of 49 samples quantified 1279 proteins. 15 samples were replicated in both assays to verify consistency between batches. In the two batches, 96.4% (161/167) and 100% (49/49) of samples passed Somalogic normalization acceptance criteria.

Throughout this paper, we use Somalogic protein marker labels, (supplementary data file S1 provides full protein names).

### Protein marker selection and predictive model building

Markers were selected based on statistical significance of differences in their abundance in the bacteria versus virus (BvV) and bacteria versus malaria or virus (BvVM) comparisons. Classifiers were developed to discriminate (1) BvV and (2) BvVM, using the 219 and 151 statistically significant (FDR<0.01) markers, respectively, and their corresponding surrogates. Using optimal subsets of N protein markers (N=5, 10, 15, 25, 50, 100) identified using genetic algorithms, 2-class Random Forest (RF) and Elastic Net (EN) models were constructed, achieving predictive results with high sensitivity and specificity with a small subset of markers (see Fig. 1; details in the “Data Analysis Pipeline” in the Supplementary Appendix).

**Fig. 1.**
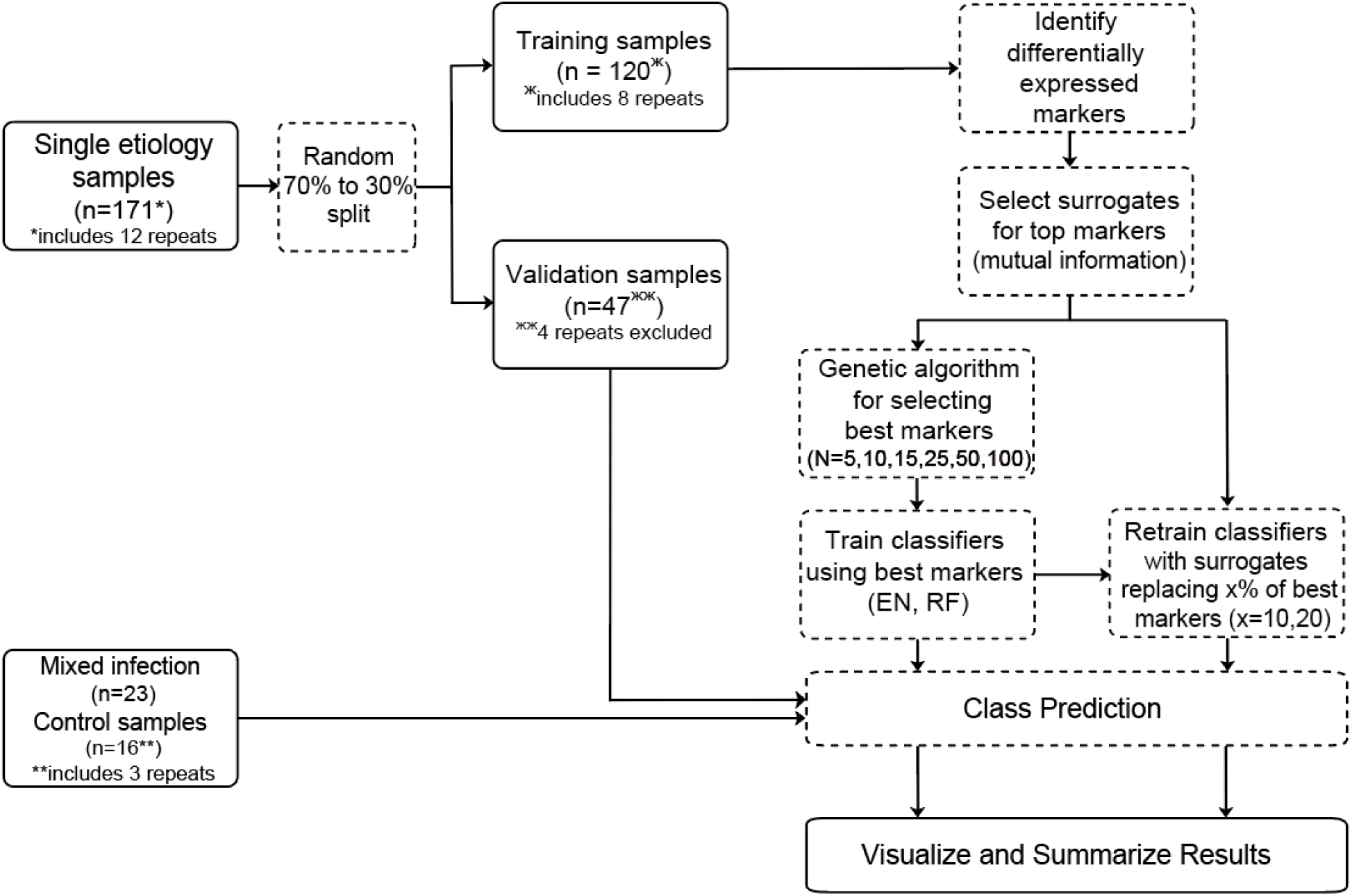
Data analysis workflow. 210 samples passed QC on the SOMAScan assay to quantify 1107 proteins. The 171 single etiology samples were classified as malaria, virus, or bacteria and included 12 repeats that were randomly split between the training and validation datasets; the 4 repeats that ended up in the validation dataset were excluded from downstream analysis. The remaining 39 samples consisted of 16 healthy community controls and 23 samples with mixed etiology. Single etiology samples were divided into a training set of 120 and a validation set of 47 samples. The training data were used for identifying differentially expressed markers between bacteria and virus, or bacteria and malaria or virus samples. Genetic algorithms were used to select the best 5, 10, 15, 25, 50, and 100 markers. Classifiers for Bacteria vs Virus (BvV) and Bacterial vs Malaria or Virus (BvVM) were trained using Random Forest (RF) and Elastic Net (EN) algorithms. Models were tuned using cross validation, and final model performance was assessed using the validation data. In order to contend with the situation where a marker is unavailable (e.g., due to difficulty in measuring the marker in a clinical setting), we determined a set of surrogate markers for each differential marker using information correlation, a criterion based on mutual information. We then assessed model performance when 10% or 20% of differential markers were substituted with their corresponding surrogates.

### Biological processes and pathways

To better understand the biological significance of the differentially expressed proteins, those statistically significant differential markers were used as input to the Metascape Gene Annotation and Analysis Resource (http://metascape.org) to query multiple ontology resources including KEGG pathway, Gene Ontology (GO) Biological Processes, Reactome Gene Sets, Canonical Pathways, and CORUM. Both three-way (Bacteria vs Malaria vs Virus) and binary (BvV) comparisons were explored (see Supplementary Appendix for details).

### Comparative marker analysis between technologies

To assess whether markers identified as indicating bacterial infection were consistent across technology platforms and between RNA and protein, extensive comparisons were made between this and two previous marker studies of the same patients. One study used RNA-sequencing and the other used multiplex bead-based protein immunoassays; both studied different but overlapping subsets of subjects within the same study population (see details in the Supplementary Appendix).

## Results

### Patient characteristics

Between July 2010 and November 2014, 576 patients were recruited as inpatients, along with 117 community controls. 195 patients under 10 years of age with acute febrile respiratory illness met the stringent inclusion criteria and were included in this analysis. To identify differentially expressed proteins between underlying etiologies, patients were characterized as having bacterial (69 patients), malaria (42 patients), viral (48 patients), or mixed (23 patients) infections. 13 healthy subjects were included as controls. The classification scheme was similar to that used previously with the same study population (see Fig. S1 for a patient classification flowchart).^9,13^ No significant differences in age, sex, weight, height, nutritional status, or duration of hospital admission were observed between bacterial, viral, and malaria sample sets. (See Table 1 and Table S1 for patient demographic and disease characteristics). Case fatality rates were high (6%) for the bacterial group, but none of the malaria cases or viral cases died. Malnutrition was highly prevalent among the three groups, and HIV prevalence was also high, although significantly higher among the bacterial group. Bacterial cases had the highest leukocyte count and respiratory rates. Malaria cases were most anemic, had the highest mean axillary temperature, and had the lowest respiratory rates. Viral cases had the lowest leukocyte count, had lower mean axillary temperature and were less anemic. Neutrophil levels were statistically higher for the bacterial etiology, but the overlap between etiologies was too great for this to serve as a classifier.

**Table 1.** Patient Demographic and Disease Characteristics at Admission.

| Features on admission (signs, symptoms, and laboratory results) | Bacteria and PCR |  |  |  |  |  | P† |
| --- | --- | --- | --- | --- | --- | --- | --- |
|  | Bacteria | No. | Malaria | No. | Virus | No. |  |
| Age (month), mean (SD) | 29.7 (29.4) | 69 | 26.3 (23.3) | 42 | 19.4 (19.9) | 48 | 0.12 |
| Female sex, n (%) | 24 (45) | 69 | 28 (67) | 42 | 23 (48) | 48 | 0.07 |
| **Clinical Examination results on arrival** |  |  |  |  |  |  |  |
| Weight (kg), mean (SD) | 10.2 (4.8) | 69 | 10.3 (4.6) | 42 | 9 (3.4) | 48 | 0.44 |
| Height (cm), mean (SD) | 80.3 (18.7) | 69 | 79.7 (18) | 42 | 74.5 (14.9) | 48 | 0.35 |
| MUAC (cm), mean (SD) | 13.5 (2) | 69 | 14.3 (2) | 40 | 14.0 (1.5) | 48 | 0.75 |
| Temperature (°C), mean (SD) | 38.2 (1.2) | 69 | 38.4 (1.4) | 42 | 37.6 (1.1) | 48 | 0.041 |
| Respiratory rate (cycles per min), mean (SD) | 60.3 (14.7) | 69 | 53.1 (8.7) | 40 | 56.7 (9.4) | 48 | 0.02 |
| **Nutritional status** |  |  |  |  |  |  |  |
| WAZ > -1 SD, n (%) | 16 (24.6) | 69 | 17 (40.5) | 42 | 24 (50.0) | 48 |  |
| WAZ -1 SD to -3 SD, n (%) (low to severe underweight) | 30 (52.2) | 69 | 18 (42.9) | 42 | 17 (35.4) | 48 |  |
| WAZ < -3 SD, n (%) (severe underweight) | 12 (23.2) | 69 | 7 (16.7) | 42 | 7 (14.6) | 48 |  |
| WAZ: Mean (SD) | -2 (1.8) | 69 | -1.5 (1.7) | 42 | -1.5 (1.7) | 48 | 0.09 |
| **Anaemia status on admission** |  |  |  |  |  |  |  |
| Hemoglobin (g/dL), mean (SD) | 8.6 (2.2) | 69 | 7.4 (2.3) | 40 | 10.0 (2.1) | 48 | <0.0001 |
| Hematocrit, mean (SD) | 26.1 (6.2) | 69 | 22.1 (7) | 40 | 29.7 (5.9) | 48 | <0.0001 |
| No anaemia (HCT > 33%), n (%) | 4 (6) | 69 | 2 (5.0) | 40 | 14 (29.2) | 48 |  |
| Mild anaemia (HCT 25 - \| 33%), n (%) | 31 (46.3) | 69 | 11 (27.5) | 40 | 25 (52.1) | 48 |  |
| Moderate anaemia (HCT 15 - \| 25%), n (%) | 32 (47.8) | 69 | 22 (55.0) | 40 | 8 (16.7) | 48 |  |
| Severe anaemia (HCT ≤ 15%), n (%) | 0 (0) | 69 | 5 (12.5) | 40 | 1 (2.1) | 48 |  |
| **Micro-biology and other laboratory results on admission** |  |  |  |  |  |  |  |
| HIV status positive, n (%) | 25 (36.2) | 69 | 3 (7.1) | 42 | 5 (10.4) | 48 | <0.0001 |
| Viral coinfections, n (%) | 32 (46) | 69 | 22 (52) | 42 | - | - | 0.56<sup>¶</sup> |
| Positive blood culture, n (%) | 29 (42.0) | 69 | 0 (0) | 42 | 0 (0) | 48 |  |
| WBC count (10<sup>3</sup>/uL), mean (SD) | 21.3 (13.1) | 69 | 14.2 (8.4) | 41 | 10.8 (2.7) | 48 | <0.0001 |
| Neutrophil granulocytes (10<sup>3</sup>/uL), mean (SD) | 13.7 (9.8) | 57 | 5.2 (3) | 32 | 4.8 (2.4) | 45 | <0.0001 |
| Plasmodium density (parasites/uL), geometric mean (SD) | 0 (0) | 69 | 5.9 (5.8) | 42 | 0 (0) | 48 |  |
| Malaria positive | 0 (0) | 69 | 42 (100) | 42 | 0 (0) | 48 | <0.0001 |
| **Chest X-Ray results** |  |  |  |  |  |  |  |
| Normal, n (%) | 9 (15.0) | 69 | 42 (100) | 42 | 27 (56.3) | 48 | <0.0001 |
| Other infiltrate/abnormality, n (%) | 7 (11.7) | 69 | 0 (0) | 42 | 21 (43.8) | 48 |  |
| Primary endpoint pneumonia, n (%) | 44 (73.3) | 69 | 0 (0) | 42 | 0 (0) | 48 |  |
| **Evolution during admission** |  |  |  |  |  |  |  |
| Length of admission (days): Median (IQR) | 4.1<br>(2 - 6.1) | 6 | 3.4<br>(2 - 5) | 4 | 3.8<br>(2.2 - 4.8) | 4 | 0.29 |
| Case fatality rate (in hospital death), n (%) | 3 (4.4) | 6 | 0 (0) | 4 | 0 (0) | 4 | 0.15 |

HCT = hematocrit; IQR = Interquartile Range; MUAC = middle upper arm circumference; n= number of patients, SD = Standard Deviation; WAZ = weight-for-age Z score, Z-score cut-off point of <-2 SD and <-3 SD is classified as low weight for age and severe undernutrition, respectively. † P-values for continuous variables were estimated through analysis of variance (Kruskal-Wallis test). P-values for categorical variables used Chi Square test. ¶ P-value of the categorical variable was estimated through Fisher's exact test.
The “bacteria” group includes blood or pleural fluid culture-positive samples, samples PCR-positive for respiratory pathogens, and samples with positive leukocytosis and a dense radiographic consolidation (endpoint pneumonia) as independently assessed by two experts. Samples that were culture or PCR positive for contaminant bacteria were excluded.

From the 195 patients, 210 peripheral blood samples (including 15 replicates, four of which were excluded from downstream analysis) were assayed for protein composition using the SOMAScan platform (see Fig. 1). Sample characteristics and designations of single (167 samples) and mixed infections with controls (39 samples) can be found in Table S2 and S3, respectively.

### Differential markers

Using the SOMAScan data, 219 and 151 differentially expressed protein markers (FDR<0.01) were identified in the BvV comparison (Table S4 A, heatmap in Fig. 2) and the BvVM comparison (Table S4 B, heatmap in Fig. S2 B), respectively. The depth and coherence of the differential signal is shown in the heatmap in Fig. 2 A. This signal is manifest only after marker selection; unsupervised clustering in the space of the entire 1107 protein panel does not reveal a clear dominant structure related to infectious etiology (Fig. S2 A). Box and whisker plots of the 100 top ranked markers are depicted in Fig. S4.

**Fig. 2.**
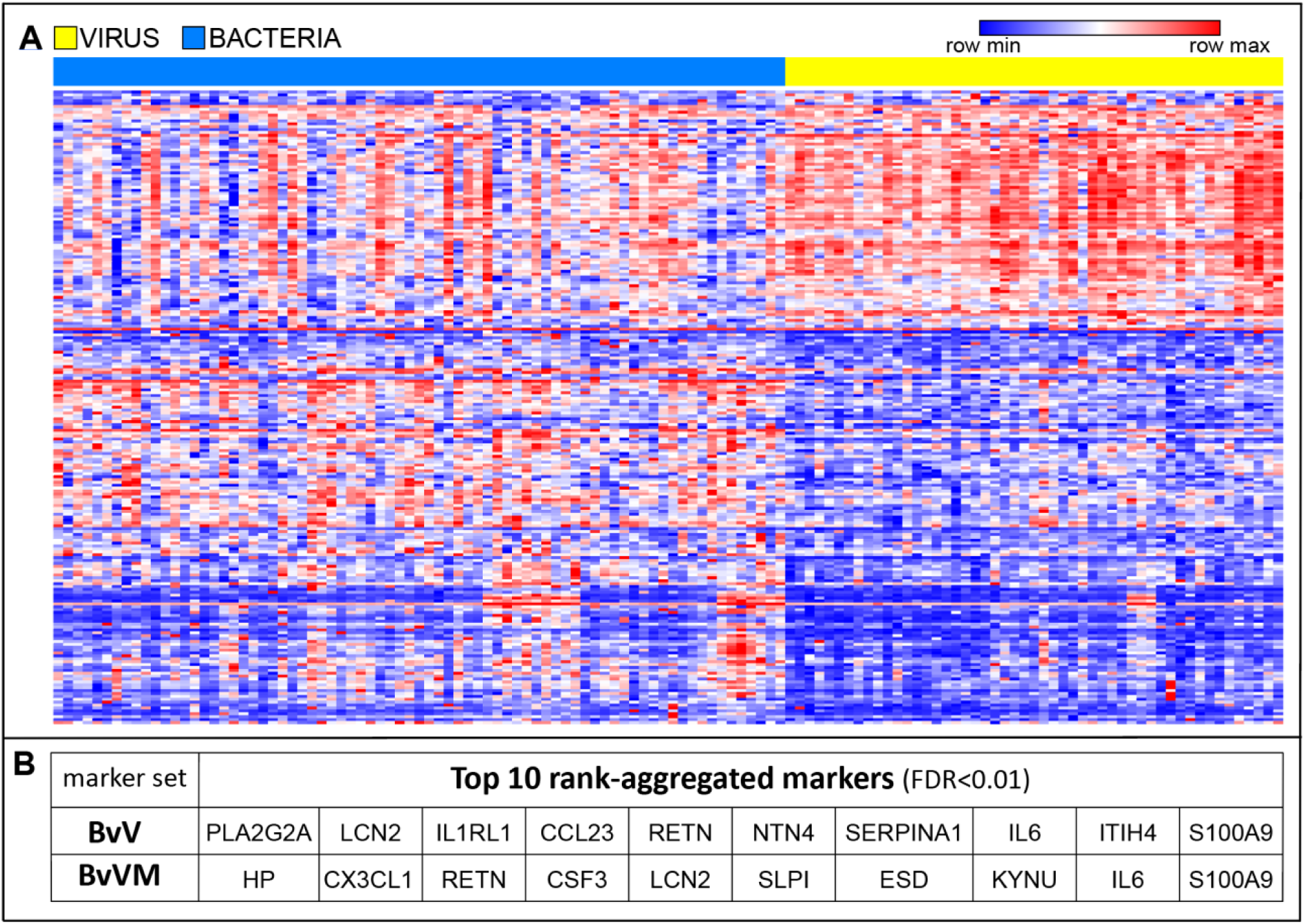
Heatmap of the Bacteria vs Virus model and top 10 rank-aggregated markers. (**A**) Hierarchically clustered heatmap of normalized SOMAscan expression values for 219 significant markers (FDR<0.01) from the SOMAScan Bacteria vs Virus (BvV) comparison in the space of all single etiology bacterial and viral samples in this study (see Fig. S10 for full resolution with details). Top track: viral (yellow) and bacterial (blue) etiology. (**B)** Top 10 rank-ordered protein markers (highest to lowest, left to right) in our BvV and BvVM marker sets. CCL23 (C-C motif chemokine 23), CSF3 (Granulocyte colony-stimulating factor), CX3CL1 (Fractalkine), ESD (S-formylglutathione hydrolase), HP (Haptoglobin), RETN (Resistin), PLA2G2A (Phospholipase A2), LCN2 (Neutrophil gelatinase-associated lipocalin), IL1RL1 (Interleukin-1 receptor-like 1), NTN4 (Netrin-4), SERPINA1 (Alpha-1-antitrypsin), IL6 (Interleukin-6), ITIH4 (Inter-alpha-trypsin inhibitor heavy chain H4), S100A9 (Protein S100-A9), KYNU (Kynureninase), SLPI (Antileukoproteinase).

### Performance of predictive diagnostic models

The chief aim of our project was to develop a protein-based biomarker panel that would distinguish bacterial from other etiologies of clinical pneumonia with accuracy that would support clinical decision-making. RF and EN models had generally similar performance, with RF models performing slightly better overall (see Table S5 C and D) and declining in performance more smoothly with fewer input markers. We therefore focused subsequent analyses on RF results.

In single etiology samples, performance of the BvV model (evaluated on the held-aside validation samples) was excellent. Sensitivity and specificity for bacterial cases using all 219 markers were 90% and 100%, respectively meeting the Foundation for Innovative New Diagnostics (FIND) proposed criteria for a diagnostic test of these characteristics.^14^ Furthermore, sensitivity and specificity remained at 90% and 85% with only 5 markers, potentially simplifying the translation to a field deployable diagnostic. Accuracy was 94% (95% CI 0.79, 0.99) and 88% (95% CI (0.71, 0.96)), with 219 and 5 markers, respectively (Table 2 A and Table S5 A).

**Table 2.**
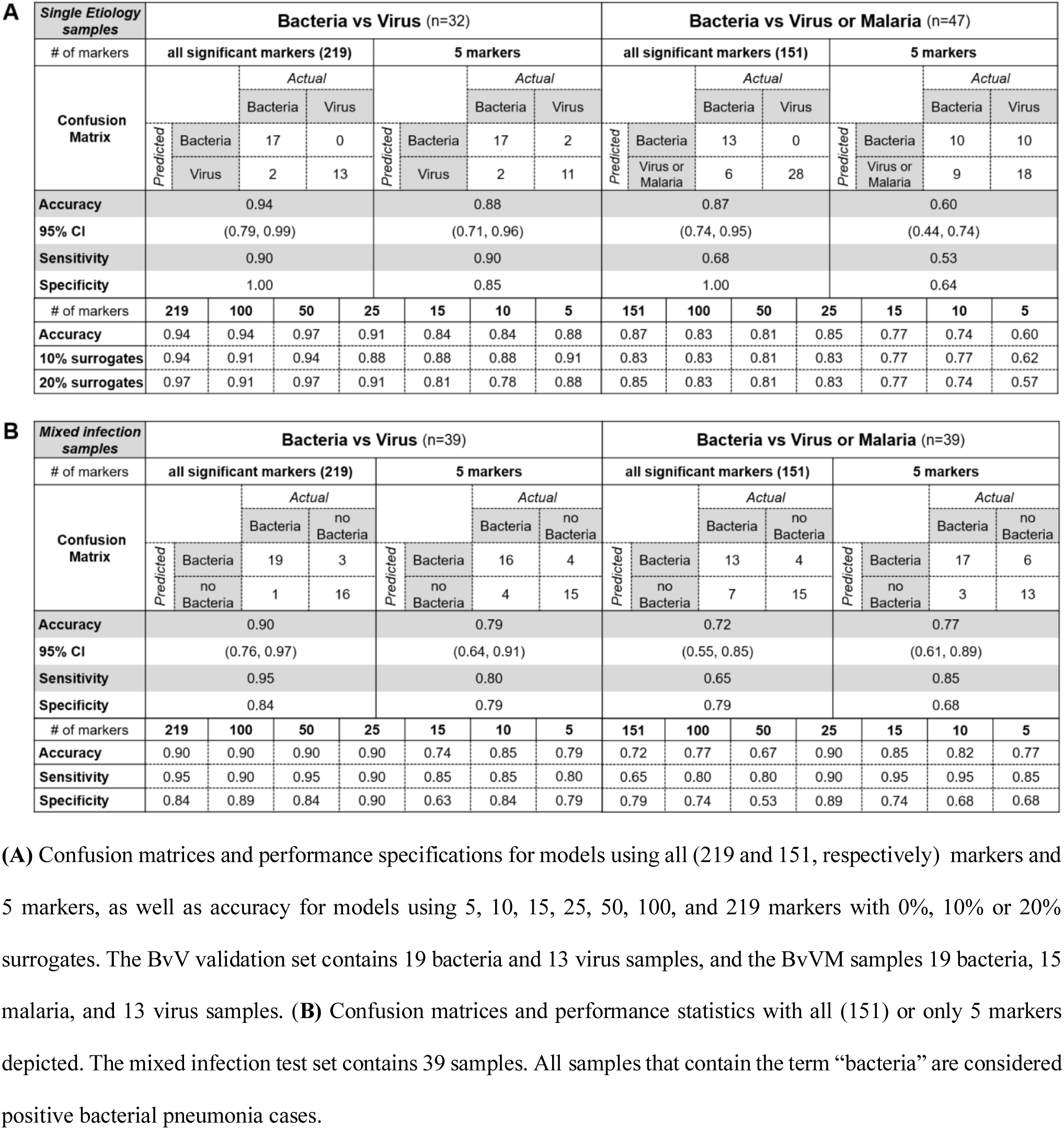
Single etiology and mixed infection validation set predictive diagnostic results.

The BvVM RF model had an accuracy of 87% (95% CI (0.74, 0.95), a specificity of 100% and a sensitivity of 68%. When decreasing the panel size to only 5 markers, accuracy decreased to 60%, specificity to 64%, and sensitivity to 53% (see Table 2 and Table S5 B).

On healthy controls and mixed infection samples, the BvV RF model performed well with 95% sensitivity, 84% specificity, and 90% accuracy (95% CI (0.76, 0.97)) (Table 2 B). The model correctly predicted the majority of bacterial infections and bacterial co-infections, successfully distinguishing these from non-bacterial infections (malaria and/or virus). Table S6 depicts BvV and BvVM RF model statistics on mixed infection samples without controls.

### Genetic algorithm-derived and surrogate markers

Marker subsets (with N ranging from 5 to 100 markers) were selected using genetic algorithms. Since the results can be nondeterministic, the method was re-run multiple times. Across all runs of the genetic algorithm, IL1RL1, HMGB1, PDCD1LG2, ROBO2, and PAPPA were the five protein markers most often selected. For the BvVM models, the most-selected markers were LTA.LTB1 (Lymphotoxin alpha2/beta1 protein), TPI1, SERPINA1, IGFBP2, and ROR1 (see supplementary data file S2 for complete marker lists).

We next assessed whether models were robust to replacement of individual markers by corresponding surrogates. This provides an index of model stability and has practical relevance when converting predictive models into diagnostics, which may require marker substitution for technical reasons. The RF (and EN) classifiers for both BvV and BvVM proved to be robust to the choice of specific markers: classifier accuracy did not significantly decline even when 20% of the markers were replaced with surrogates (Fig. S5).

### Biological processes and pathway analysis

To gain insight into the biology underlying the markers of bacterial and viral infection, multiple databases were queried for functional and pathway annotations. Terms significantly enriched in the bacterial or viral pneumonia marker sets were automatically clustered into non-redundant groups (details in methods). Marker support for terms is shown in Fig. 3 A and the top twenty clusters in Fig. 3 B. Individual terms (and therefore clusters) could be supported by both bacterial and viral markers. Most clusters had support from both etiologies, but a subset (blue or red circles in Fig. 3) was strongly associated with a single etiology. Two GO clusters, *chemotaxis* and *regulation of neurogenesis*, were driven almost exclusively by viral markers, while *response to bacterium* (our top ranked GO term with 39 gene hits), *regulated exocytosis, antimicrobial humoral response, positive regulation of response to external stimulus*, and *signaling by interleukins* were driven almost exclusively by bacterial markers.

**Fig. 3.**
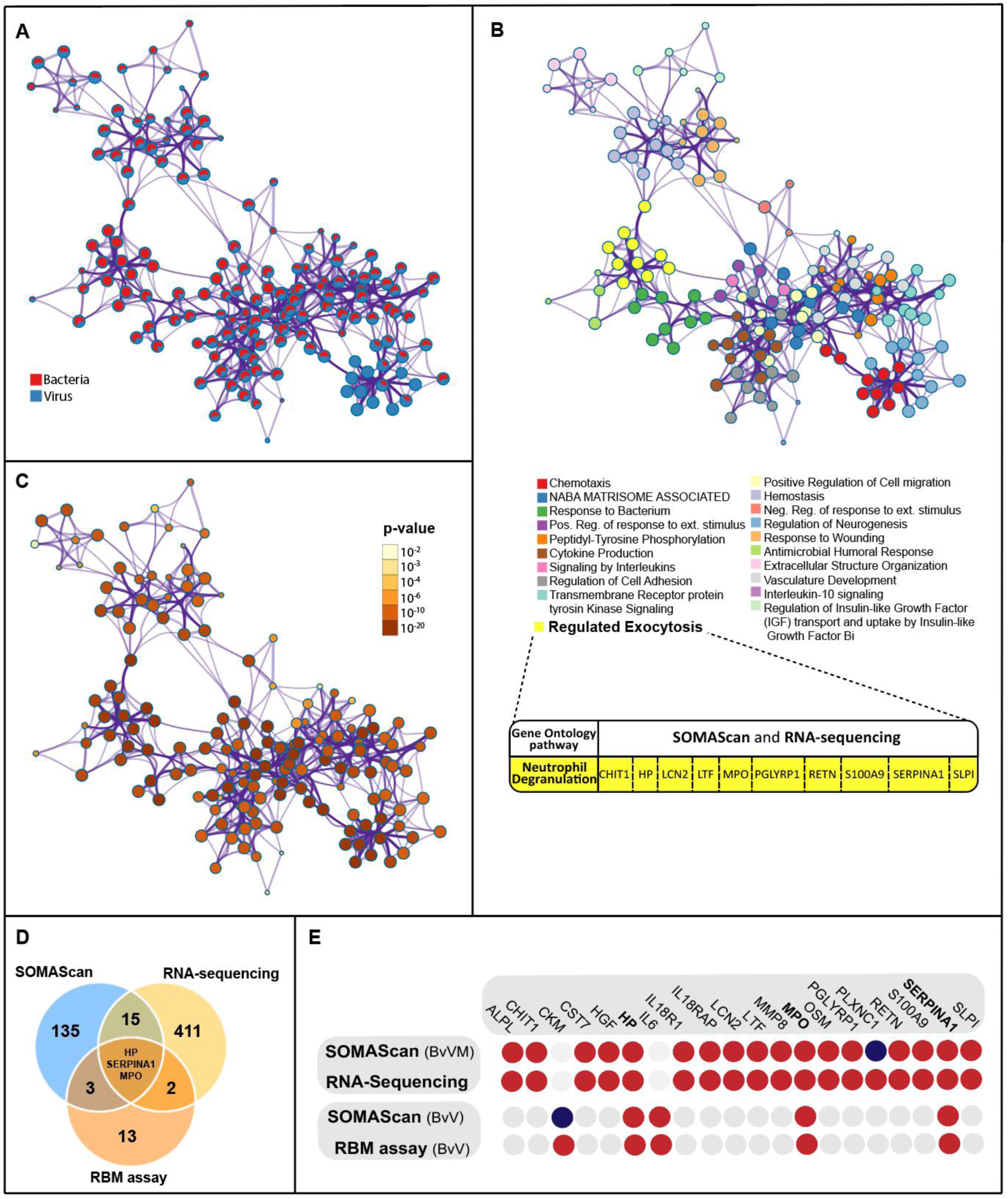
Pathways and gene enrichment analysis with differential markers shared between this study, RNA-sequencing, and RBM multiplex assay studies with the same study population. **(A,B,C)** Clustered terms enriched in our bacteria vs virus 2-class comparison. Each node represents one term describing a biological process or pathway. Edges connect similar terms (similarity score (κ) > 0.3); the thickness of the edge represents the similarity score. Each term is represented by a circle node, where the size is proportional to the number of input markers. The underlying file can be found as an additional supplementary file (“Cytoscape BvV network”). **(A)** Distribution of support for each node from bacterial (red) and viral (blue) markers, i.e. each pie sector is proportional to the number of hits that originated from a particular marker list. **(B)** Nodes colored by their membership in one of the top 20 clusters. Each cluster is named for the term (node) with the best p-value. Inset table: neutrophil degranulation, considered as a sub-pathway of regulated exocytosis, was detected as the major biological GO pathway shared between the BvVM marker set of this study and RNA-sequencing data. Of the 18 bacterial markers overlapping between the studies, 10 markers are directly involved in neutrophil degranulation (see Fig. 3E for all 18 markers). **(C)** Bacteria vs Virus marker set with nodes colored by p-value. The darker the color, the more statistically significant the node (see legend for p-value ranges). **(D and E)** RBM and SOMAScan protein aliases were converted into their gene names to compare markers between studies. **(D)** Overlap of selected marker sets: SOMAScan (BvVM, n=156), RNA-Sequencing (BvVM, n=431), and RBM immunoassay (BvV and BvM, n=21). **(E)** Two direct comparisons of marker sets derived through the same approach (bacteria vs virus (BvV) and bacteria vs virus or malaria (BvVM)); filled circles indicate a marker identified in the specified analysis. Markers that overlapped in the two direct comparisons are depicted by filled circles, but not between the four individual marker sets. The color indicates the direction of expression change. Red: upregulation in bacterial samples; dark blue: downregulation. Light grey: the marker was not detected or not included in at least one of the two marker sets. Haptoglobin (HP), Myeloperoxidase (MPO), Alpha1-Antitrypsin (SERPINA1)

Neutrophil-related biological processes emerged as a key biological theme associated with bacterial infection. In particular, the r*egulated exocytosis* GO cluster (34 gene hits) is mostly comprised of neutrophil- or leukocyte-related terms. Within the top 36 GO clusters (out of 1388 total clusters, ranked by p value), six highly significant clusters that consist of 14 to 26 gene hits each were identified as neutrophil processes (*migration, mediated-immunity, activation, degranulation, activation involved in immune response*, and *chemotaxis*). Notably, no other cell type or subpopulation besides neutrophils appeared within the first 243 rank-ordered GO clusters. The *neutrophil degranulation* cluster was particularly prominent in markers that were identified by both SOMAScan and RNA-sequencing; it contained 10 of the 24 markers that emerged from that cross-platform comparison (Fig. 3 B, D, E).

### Comparisons between datasets and technologies

To assess the consistency of the results, we compared gene marker sets from similar marker-focused studies of the same population using different technologies. First, the current BvVM marker set was compared with markers found in our previously published RNA-sequencing approach.^9^ Of the 1107 proteins included in our SOMAScan assay, 78 were represented by genes from the set of 600 significant differentially expressed markers in the RNA-sequencing analysis (of ∼12,000 expressed genes) (Data file S1 D). Twenty-five of these 78 genes (corresponding to 24 proteins) proved to be statistically significant markers in our comparison (Fig. S6).

In the RNA data, 18 of those 24 proteins were markers for bacterial infection and 6 were markers for malaria infection. A heatmap of these markers highlights the strong class distinctions (Fig. S7). Haptoglobin (HP) is markedly down and hemoglobin up in malaria samples, but the majority of markers are elevated in bacterial samples (Fig. 3E, and see Fig. S6 and Fig. S8 for details on the malaria markers). When we used the SOMAScan data for these 24 markers to build RF and EN models, they performed similarly to 25 protein marker models optimized by the genetic algorithm (Table S5 E and F), suggesting that those 24 markers would also be good candidate markers for a diagnostic assay.

We also compared the SOMAScan marker sets with findings from a previous protein-based immunoassay (the RBM multiplex immunoassay).^13^ Five markers were identified as differential markers for bacterial pneumonia in both datasets: CKM, HP, IL6, MPO, and SERPINA1 (Fig. 3E, Fig. S6). Three markers, Haptoglobin (HP), Myeloperoxidase (MPO), and Alpha1-Antitrypsin (SERPINA1), were identified as significant markers in all three studies (SOMAScan, multiplex immunoassay, and RNA-sequencing) despite the very different methodologies employed (Venn diagram in Fig. 3D). Two markers appear in both the SOMAScan and multiplex immunoassay data as likely markers for malaria infection, VCAM1 and APCS.^13^

## Discussion

Here we present diagnostic models based on aptamer-derived blood protein signatures that accurately discriminate bacterial from viral infections of pediatric febrile respiratory illness with as few as 5 protein markers (94% accuracy, 90% sensitivity, 85% specificity). These models meet or exceed the performance guidelines proposed by a FIND-sponsored expert consensus document for a diagnostic test of bacterial pneumonia.^14^ Accurate discrimination of bacterial infection from both viral and malaria etiologies was achieved with 25 markers.

Because the BvV model was highly predictive, we investigated the underlying marker proteins for information about the inflammatory processes that typify bacterial and viral infections. Gene enrichment and pathway analyses showed that the processes in bacterial infections were dominated by neutrophils. The central role of neutrophil biology in the host response to bacteria is highlighted by the consistency of that signal across prior studies at both the RNA and protein level, with an overlap of 18 bacterial markers despite differences in both the assay platforms and the model-building approaches used (Fig. 3 E). Reinforcing this observation, a cross-platform 24 marker set that we identified (which was highly enriched for neutrophil-associated proteins) proved to be equally effective in differentiating bacteria from other causes of pediatric pneumonia (Table S5 F). The neutrophil degranulation pathway was particularly enriched in the cross-platform marker set. Ten of the 18 bacterial markers in the cross-platform analysis were associated with bacterial airway inflammation, modifying, mitigating or augmenting neutrophil immunological responses. For example, SERPINA1 and SLPI are both protease inhibitors regulating neutrophil elastase activity.^19–21^

The overarching objective of this work was to develop protein-based predictors that could be ported to a field-deployable device for discriminating bacterial from non-bacterial pneumonia. While larger validation studies are needed, this study provides strong evidence that a blood-based protein panel of limited size can achieve the sensitivity and specificity required to guide clinical decisions regarding antibiotic therapy in febrile children with respiratory distress. It also lays the groundwork for future development of a point of care test by identifying biologically plausible sets of markers that could serve as its basis, particularly considering that some of these markers (haptoglobin, SERPINA1, MPO etc.) are relatively simple to measure. We have also identified surrogate proteins that can be exchanged for markers in our models without loss of accuracy, allowing flexibility in developing a diagnostic test. Though optimized for single etiology samples, our models performed well in mixed infections, which represent the true natural complexity of febrile respiratory illness. Importantly, these markers seem to discriminate appropriately, even in the context of a high underlying malnutrition or HIV prevalence, such as the one in Manhiça, where the study was conducted.^22,23^ This is a significant benchmark, as a predictor must be effective across the spectrum of real-life clinical scenarios. Finally, our study has also provided insights into the biology of host response as reflected in discriminant marker proteins. These observations may inform marker selection in future prospective studies, and together with our specific models and markers may facilitate the development of the optimized point-of-care tests that are needed to change future clinical practice, particularly for those settings where associated case-fatality rates for common infections remain high and diagnostic tools scarce.

## Acknowledgments

We gratefully acknowledge the crucial commitments of our colleagues Godfrey Allan Otieno, Jacob Silterra, Katherine Almendinger, Karsten Krug, Roger Wiegand, and Rushdy Ahmad to the overall success of this clinical study, especially in the early phases.

## Data and materials availability

The original Somalogic data files along with matching de-identified clinical data are available through collaboration with Dr. Quique Bassat. The code is available as .zip file in the supplementary material and includes (i) code used for the study (ii) datasets and intermediate tables and (iii) results. README.txt file is available.

