## Supplemental Figures for "Biomarkers to distinguish bacterial from viral pediatric clinical pneumonia in a malaria endemic setting"

This appendix has been provided by the authors to give readers additional information about their work.

**Methods** 3
Study population 3
Sample Collection 3
Etiologic Diagnosis Classification 4
Ethics 4
Data analysis pipeline 5
Marker selection and model building 5
Differential marker selection 5
Identification of surrogate markers 6
Selection of the most predictive subset of markers through genetic algorithms 6
Evaluating the accuracy of a clinical signature: ALMANACH 7
Biological processes and pathways 7
**Results** 10
Study population 10
Differentially expressed markers 10
**Discussion** 13
**Figures** 17
Fig. S1 Single etiology patient classification 17
Fig. S2 Heatmaps of unsupervised hierarchical clustering of all 1107 proteins and BvVM markers (n=151) 18
Fig. S3 Protein marker overlap of the 30 highest ranking markers. 20
Fig. S4 Box and whisker plots of the 100 Top markers. 21
Fig. S5 Model accuracy when switching out surrogates for comparisons of interests 23
Fig. S6 RBM multiplexed Immunoassay, RNA-sequencing and SOMAscan assay comparisons with Venn diagrams and marker lists 25
Fig. S7 Heatmap of all 24 overlapping SOMAScan (BvVM) and RNA-sequencing markers. 29
Fig. S8 Malarial markers shared between SOMAScan and RNA-sequencing. 30
Fig. S9 Enrichment Heatmap of the three-way comparison (Bacteria vs Malaria vs Virus). 31
Fig. S10 Heatmap of the Bacteria vs Virus model. (A) Hierarchically clustered heatmap of normalized SOMAscan expression values for 219 significant markers (FDR<0.01) from the SOMAScan Bacteria vs Virus (BvV) comparison in the space of all single etiology bacterial and viral samples in this study. 32
**Tables** 33

Table S1 Patient demographic and disease characteristics at admission for Bacterial and PCR Bacterial samples. 33

Table S2 Single etiology sample designations and distributions for training (n=120) and validation sets (n=47). 35

Table S3 Mixed infection sample designation sample distribution. 36

Table S4 List of markers for Bacteria vs Virus (n=219) and Bacteria vs Virus or Malaria (n=151). 38

Table S5 Single Etiology validation statistics for BvV and BvVM marker sets (A to D), and SOMAScan assay and RNA-Sequencing overlapping marker set (n=24) for RF and EN models (E and F) 55

Table S6 Mixed infection set without controls (n=23) (Random Forest) 60

Table S7 ALgorithm for MANagement of Childhood illnesses (ALMANACH). 62

**References** **64**

#### Methods

##### Study population

Exclusion criteria are identical to those in Valim et. al., 2016[1,2]: use of antimalarial drugs in the preceding 2 weeks, established or suspected pulmonary tuberculosis (based on a history of cough of more than 2 weeks' duration or history of direct contact with a patient with active tuberculosis), ongoing participation in conflicting studies, and severe hypoxemia (oxyhemoglobin saturation ≤85% or less) to reduce the likelihood of *Pneumocystis jirovecii* infection. Exclusion of suspected infections by *Mycobacterium tuberculosis* and *Pneumocystis jirovecii* aimed to improve the specificity of case definitions.

Healthy community controls were matched to cases by sex, age, and neighborhood and identified through a demographic surveillance system, which allowed for selection of a random sample of children from the same age distribution and geographic area as the study patients. Control patients were examined and excluded if they presented with fever or respiratory symptoms, if they had taken any medication in the previous 30 days, or if they had a positive malaria microscopy exam.

Operators blinded to diagnosis and clinical status processed all laboratory tests and samples. All clinical, laboratory, and radiologic data were used to classify all patients enrolled in the study as having bacterial, viral, or malaria infection, into either single or mixed etiologies (both used in this study).[2]

##### Sample Collection

Before the initiation of antimicrobial treatment, a nasopharyngeal aspirate (NPA) was obtained to determine viral respiratory infections through a real-time polymerase chain reaction (RT-PCR) test. Additionally, venous blood was collected for blood culture, bacterial PCR for respiratory pathogens (referred to as “PCR Bacteria samples” if positive), human immunodeficiency virus (HIV) testing, malaria detection by smear microscopy, and SOMAScan protein assay. Pleural fluid, when available, was also submitted for culture. Coagulase-negative staphylococci, *Bacillus* species, or *Micrococci* were considered contaminants. Bacterial RT-PCR was performed for *Haemophilus influenzae* (HiB) and *Streptococcus pneumoniae* (pneumococcus), the two most common causes of bacterial pneumonia in this pediatric population.[2] Detection of the *bexA* gene was considered diagnostic for HiB[3], while the *ply* and *wzg* genes identified pneumococcus if confirmed by at least one positive hit in two independent PCRs. [3,4] In 32 out of 58 tested samples both *ply* and *wzg* were positive.

**Etiologic Diagnosis Classification**

Single etiology cases generally required absence of evidence of other etiologies, and definitions were mutually exclusive. However, the bacteria and malaria classifications allowed concurrent viral infections in the NPA. A comprehensive flow chart for patient single etiology classification can be found in Fig. S1.

**Ethics**

This study was reviewed and approved by the Mozambique National Bioethics committee (reference No. 262/CNBS/10), the Institutional Review Board of the Broad Institute/MIT Committee on the Use of Humans as Experimental Subjects (reference No. 1003003765), and the Hospital Clinic’s Ethics committee in Barcelona, Spain (reference No. 2010/5590). A signed informed consent of parent or guardian was obtained prior to study participation. The presence of a witness was required in case of illiterate guardians.

#### Data analysis pipeline

##### Marker selection and model building

Single etiology samples (n=167) were randomly divided into a training set (70% of samples, n=120) for model building and an independent validation set (30% of samples, n=47, after excluding 4 repeated samples to prevent information leak from the training data) (Table S2). Markers selected in the training set were used to train models that were evaluated in the validation set. The same models were also tested in a separate sample set comprising the 23 mixed infection samples, 13 healthy controls, and 3 replicates (Table S3). Fig. 1 shows an overview of the data analysis workflow for single etiology and mixed infection samples. Because the critical focus was on identifying bacterial samples to guide antibiotic therapy, markers were selected and evaluated by two different 2-class comparisons of diagnosis groups: bacteria versus virus (BvV) and bacteria versus malaria or virus (BvVM).

##### Differential marker selection

Markers were first selected based on statistical significance of differences in their distributions in the BvV and BvVM comparisons estimated through SAM (Significance Analysis of Microarrays), with p-values corrected for multiple testing through the Benjamini-Hochberg FDR (False Discovery Rate) approach.[5] SAM analysis and statistical significance determination were performed in R using the samr library. Markers that were statistically significant ($FDR < 0.01$) were then used as input to train multiple classifiers including partial least squares, partitioning around medioids, random forests (RF), and elastic nets (EN). The importance of each input marker to overall classifier accuracy was determined using variable importance metrics from each of the

4 classifiers. The ranking of markers based on SAM and classifier variable importance were combined into a single final rank using robust rank aggregation.[6] Classifier training and variable importance determination were coded using the caret library in R.[7]

**Identification of surrogate markers**

We sought surrogates for all markers with FDR<0.01 by calculating the mutual information-based information correlation (IC) between each marker and all other proteins in the dataset. Any protein with IC>0.75 was considered a surrogate for the corresponding marker [8]. This analysis provided important latitude for final assay construction, as a surrogate marker could replace its companion candidate marker in a diagnostic test to address challenges such as reagent availability or inadequate analytical performance.

**Selection of the most predictive subset of markers through genetic algorithms**

As assays with fewer markers are generally easier to deploy, we aimed to identify subsets of markers that efficiently captured our classifications of interest -- distinguishing bacteria from virus, and bacteria from virus or malaria. The goals were to identify a small number of markers that effectively classified the samples, to assess the effect of adding more markers on model performance, and to compare reduced marker models to those trained using all statistically significant markers. To this end, we applied genetic algorithms to find the best subset of N markers (with N = 5, 10, 15, 25, 50, 100) for RF and EN classifiers. The genetic algorithm was implemented in R using the rgenoud library.[9] The BvV or BvVM protein marker sets and their surrogates (Data file S3) were the input markers to the non-deterministic genetic algorithm. The fitness function (or objective function) for the genetic algorithm was based on the accuracy of RF or EN models trained using the marker subset under consideration, and evaluated on a held-aside subset of the input training dataset. As the marker subsets evolved, the fitness function was maximized to obtain the best subset of N markers.

Once the genetic algorithm identified the best markers for each N, repeated 10 times with random restarts, we ranked the markers based on the cumulative frequency with which they were identified in all runs. We selected the N most frequently identified markers based on this ranking, and used them to refit a random forest and elastic net model using the entire training dataset. The models were then applied to the validation set of subjects, and the predicted class of each sample based on each classifier was recorded and the misclassification rate of each classifier determined.

To contend with potential situations in which a particular selected marker could not be included in an assay (for instance, due to reagent availability, or difficulty measuring the marker in a clinical setting), we assessed model performance when randomly selected subsets of 10% or 20% of differential markers were replaced by their corresponding best surrogate markers.

**Evaluating the accuracy of a clinical signature: ALMANACH**

To compare the effectiveness of clinical parameters alone for assigning pathogenic etiology of febrile respiratory distress in our sample set, we used available clinical variables from our sample population in the context of the ALMANACH (ALgorithm for MANagement of Childhood illnesses)[10] algorithm to differentiate between bacterial and viral pneumonia (see Table S7).

**Biological processes and pathways**

Marker lists (FDR<0.01) derived from SAM-based marker selection were mapped to a single gene name for each marker. The Metascape Gene Annotation and Analysis Resource was used to query multiple ontology resources. Binary markers were assigned fold change values; proteins higher in bacterial or viral samples were assigned to Bacteria or Virus, respectively. Markers in the three-way comparison were assigned contrast scores, which represent standardized differences between the class mean and the overall mean. Proteins with positive contrast scores in a class were considered to be enriched in that class. Therefore, in the binary comparison proteins were uniquely assigned to one class, whereas in the three-way comparison proteins could be shared by two (but not all three) classes. The resulting networks can be probed in greater detail to identify individual node terms, associated Benjamini-Hochberg corrected q-values, etc. by importing the accompanying .cys file into Cytoscape (supplementary data file "Cytoscape B vs M vs V 3-class network").

**Verifying Consistency of Our Selected Markers - Comparative marker analysis between technologies**

We first derived differential marker gene lists from markers identified in: a) the BvV and BvVM comparisons within SOMAScan protein assay data (Data file S2); and b) 2-class single marker comparisons of the groups Bacteria vs Virus (BvV), Bacteria vs Malaria, and Virus vs Malaria within a multiplex immunoassay.[2] Since our prior RNA sequencing study[1] included the same Bacteria vs Virus and Malaria (BvVM) 2-class comparison that we used in the SOMAScan analysis, the resulting gene marker sets could be used without modification in these comparisons. Our 151 SOMAScan BvVM markers were mapped to 156 genes and compared with the RNA-sequencing BvVM 431 gene marker set (see overlap depiction in Fig. 4), as well as with other sets from that analysis (Fig. S6). The combined significant markers from our RNA-sequencing and multiplex immunoassay were compared to the SOMAScan BvVM (n=156 genes) or to the BvV gene marker sets (n=226). Most of these comparisons between marker sets and resulting Venn diagrams can be found in Fig. S6.

### Results

##### Study population

No significant differences in patient demographic and disease characteristics were identified between Bacteria and PCR Bacteria samples (Table 1), except neutrophil levels, suggesting that positivity in RT-PCR for bacteria allowed inclusion of a wider spectrum of disease severity in the bacteria group.

##### Differentially expressed markers

We performed marker selection to identify proteins that were differentially abundant between bacterial and viral samples (BvV), and bacterial vs the combined viral and malaria samples (BvVM). Statistically significant markers were used as input to train our models. The markers were ranked by a combination of their differential protein expression and contribution in accurately classifying samples into appropriate subgroups.

The highest ranking 10 proteins (Fig. 2B) were (gene names in parentheses): Phospholipase A2 (*PLA2G2A*), Lipocalin 2 (*LCN2*, also known as neutrophil gelatinase-associated lipocalin or NGAL[11]), C-C motif chemokine 23 (*CCL23*), Interleukin-1 receptor-like 1 (*ILRL1*), Myeloperoxidase (*MPO*), Fractalkine (*CX3CL1*), Resistin (*RETN*), Interleukin-6 (*IL6*), Netrin-4 (*NTN4*), and Kallistatin (*SERPINA4*). The 10 highest scoring proteins were Haptoglobin (*HP*), Fractalkine, Resistin, Granulocyte colony-stimulating factor (*CSF3*), *LCN2*, Secretory Leukocyte Protease Inhibitor (*SLPI*), S-formylglutathione hydrolase (*ESD*), Kynureninase (*KYNU*), Interleukin-6, and Protein S100-A9 (*S100A9*).

Seventeen of the 30 top ranking protein markers were common to both the BvV and BvVM comparisons (Fig. S3). Fractalkine, Interleukin-6, and Resistin were amongst the top 10 markers for both models. Lipocalin 2 ranked within the top 5 markers in both models. Surrogate markers for Lipocalin 2 are Myeloperoxidase and Resistin (Data File S3). Expression values for the 100 top-ranking protein markers for each comparison are depicted in Fig. S4.

**Clinical algorithms underperform protein-based predictive models in differentiating bacterial from other causes of pediatric respiratory distress**

To test the effectiveness of clinical parameters alone for assigning pathogenic etiology of pediatric respiratory distress in our sample set, we used available clinical variables from our sample population in the context of the ALMANACH (ALgorithm for MANagement of Childhood illnesses [10] algorithm for "Assessment of FEBRILE Childhood Illness" and sub-algorithm on "Cough-Related diagnoses". As our focus was on guiding antibiotic administration, we used the two clinical variables that led to a directive for antibiotics: respiratory rate above 50 breaths per minute in a calm child and "Chest indrawing or stridor in a calm child", indicative in the algorithm of "pneumonia" or "*severe* pneumonia", respectively. In our data, stridor overlapped completely with indrawing or elevated respiratory rate and was therefore not used. The implementation of the ALMANACH algorithm for this dataset was:

> *Given fever and cough or difficulty breathing (present in all of our subjects): If chest indrawing is present or sustained respiratory rate is greater than 50 breaths per minute, assign a "bacteria" etiology; otherwise assign an "other" etiology*

*(where "other" can be either "malaria or virus" or "virus" depending on the specific comparison).*

Both two-way (BvV) and three-way (BvMV) comparisons were included, and predictive accuracy in the relevant test set was compared to the protein-based predictive models. Resulting confusion matrices, statistics, and additional details can be found in Table S7.

The performance accuracy for both sample sets was poor (0.55 to 0.74), worse than our Somalogic models, and partially below the No Information Rate (NIR) for the BvVM sample set. Sensitivity was high (93%) in the BvV sample set in which a presumptive bacterial infection (pneumonia or severe pneumonia) was clinically assessed by the presence of either positive chest indrawing or a respiratory rate of over 50/min, but specificity was unacceptably low at 23%. Classifiers based on these two clinical variables separately had higher specificity and lower sensitivity than the combination. While chest indrawing as a single predictor for pneumonia performed best overall amongst the clinically-based models (accuracy 0.67, NIR 0.52, sensitivity 86%, specificity 46%), none of the models approached the specifications needed for a diagnostic assay for bacterial pneumonia and none competed with the Somalogic-based models.

#### Discussion

Since clinical parameters continue to be used to attempt to distinguish causes of fever in resource-limited settings, as for instance in the ALMANACH implementation[10], comparison of their performance to that of our predictive models is important. Taking an ALMANACH recommendation for antibiotic administration as a functional surrogate for bacterial etiology, we compared the accuracy of ALMANACH predictions to those of our protein-based models. The clinical models were uniformly inferior to our molecular predictors, with a particularly dramatic loss of specificity (see Table S7 for details). We did not attempt to incorporate clinical parameters into our protein-marker based models for disease etiology, as our goal was a test for the presence or absence of bacterial cause of febrile respiratory illness that did not rely on either resource-intensive testing (such as blood culture or radiography) or subjective clinical assessments that are often unreliable in community health settings. Furthermore, laboratory parameters (such as leukocyte count) contributed to our definition of single etiology samples, so their inclusion would generate biased models. Of note, neutrophil counts have been shown in previous studies not to differentiate disease etiologies in febrile children.[12,13]

Neutrophils, part of the innate immune system [14,15], migrate to sites of inflammation to help to eliminate bacteria, and play a central role in the pathogenesis of many respiratory diseases.[16,17] Though neutrophil infiltration in the lower respiratory tract is a hallmark of bacterial infection [17,18], their role in the pathological mechanisms of disease is not fully understood.

Evidence for other biologically important pathways can also be seen in these data, but the details are beyond the scope of this paper. For example, “interleukin-4 and interleukin-13” signaling was enriched in markers from both the SOMAScan and RNA-sequencing studies. A three-way comparison (Bacterial vs Virus vs Malaria) revealed additional, largely expected, pathways involved in each of the three disease etiologies (Fig. S9). Hemostasis signals were increased in all types of infection; thrombosis is a well-known and dangerous complication especially of bacterial and viral sepsis.[19] The appearance of a matrisome-associated protein signal in plasma, especially in bacterial infections, suggests ECM degradation. Invasive pathogens breach the basal lamina, bind to ECM and can degrade interstitial ECM proteins using proteases.[20]

Other ontologies were more strongly enriched in one or two of the categories. Especially potent cytokine production and inflammatory response signatures were shared by malaria and bacterial infections. Hemozoin and other toxic factors from the parasites may stimulate monocytes and macrophages in a fashion comparable to that of bacterial endotoxin, though other mechanisms of inflammatory induction may differ.[21] High levels of predominantly proinflammatory cytokines are responsible for the high fevers associated with blood-stage malaria infection, while elevations of both pro- and anti-inflammatory cytokines are associated with severe bacterial pneumonia and attendant respiratory failure and multi-system organ dysfunction that can accompany bacterial pneumonia.[22,23] Regulation of cell adhesion was strongly upregulated in malaria. These gene products have been suggested to contribute to the virulence of P. falciparum relative to other human malaria parasites [24], and to the clinical presentation of severe relative to asymptomatic malaria.[25] The potential significance of upregulation of Cell Adhesion Molecules (CAMs) in Virus is twofold; CAMs are often upregulated in the host defense against microorganisms[26], but can also function as viral receptors and promote infection and viral internalization.[27] Although antibodies are the principal adaptive response against both extracellular viral and bacterial pathogens, humoral immune response was most strongly represented in bacteria samples. Regulation of endopeptidase activity, which plays a critical regulatory role in both immune activation and inactivation[28] was also especially robust in bacteria samples.

Our study shares with other biomarker studies in clinical pneumonia the difficulty of confidently assigning a bacterial or viral etiology to the clinical syndrome.[29,30] Bacterial culture is insensitive and may be associated with severity of disease; therefore a bacterial class definition based on culture may bias results towards severity rather than etiology. To circumvent this problem and generate a more representative bacterial class, we included samples positive by blood-based RT-PCR for bacterial respiratory pathogens, and followed other authors in using “probable diagnosis” criteria based on expert panel review of clinical, radiologic and laboratory information [31]. Though such “probable bacteria” samples were in the minority, this greater inclusivity could result in false positive detection of bacterial etiology and misclassification bias. However, these biases would generally lead to underestimating differences across groups and not to false discovery of markers. There are additional limitations. Genetic algorithms (GAs) used for marker selection are not deterministic, so repeated runs are likely to reach different solutions, especially when the candidate marker population is large.[9,32] Relatedly, GAs do not exhaustively explore the feature landscape, so while they should achieve well-performing models, it cannot be claimed that their output results in the best possible predictor. These factors contribute to the observation that the 24 marker model based on overlapping features from multiple datasets with partially different markers (see Table S6 E and F) has equivalent performance to the GA-optimized 25 marker model (see Table S6 A to E, Table 1 and 2). Our markers were selected and models trained on single etiology samples, which importantly allowed virus-positive NPAs in the bacterial and malarial classes. While a bacterial signature could still be detected in other mixed infection samples, it is likely that performance in those mixed infections could be further enhanced by specifically training on that population. Finally, the generalizability of the model would be better assessed in the context of a larger, more ethnically and geographically diverse sample population drawn from areas with and without endemic malaria.

### Figures

#### Fig. S1 Single etiology patient classification

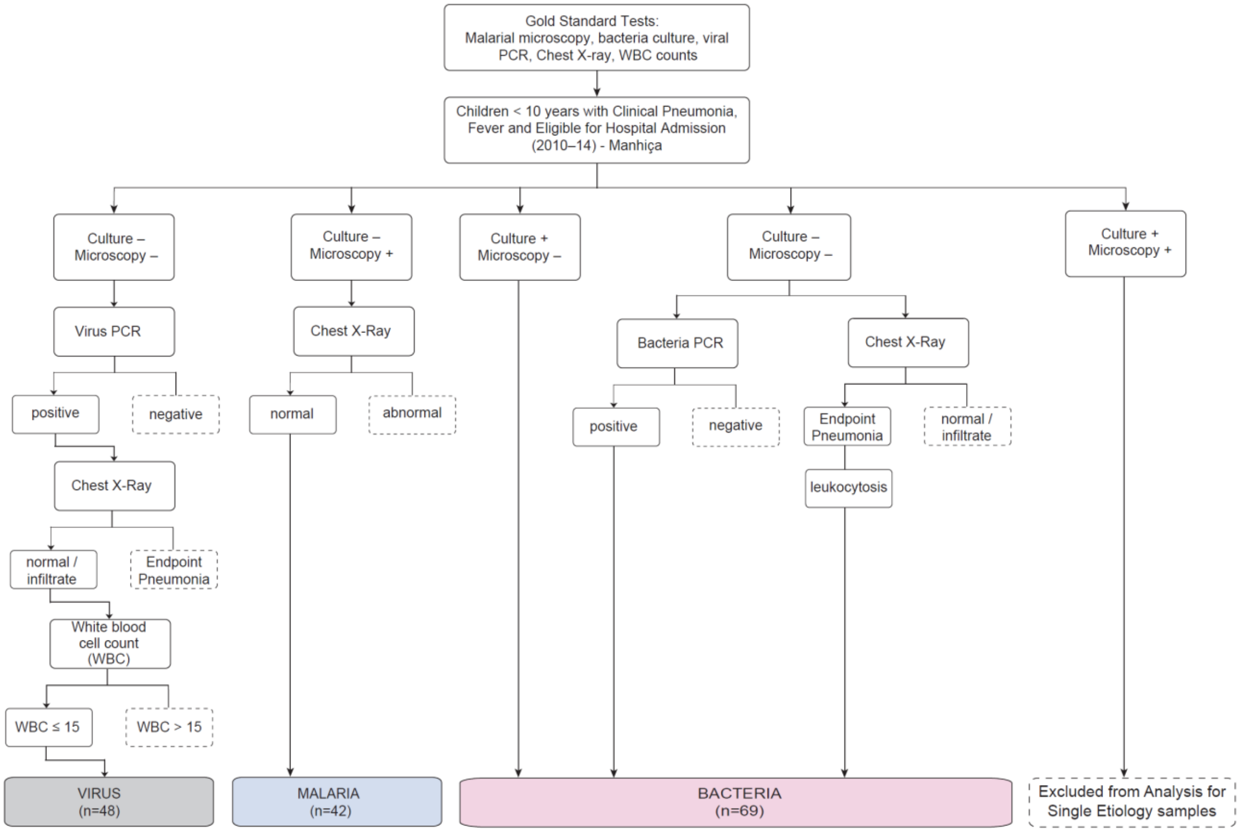

Patient classification into three single etiology groups (bacteria, virus, and malaria). Negative tests included those that had contaminant bacteria, those that were missing results, and those that did not meet the microscopic parasitemia threshold. Chest X-ray results were classified as (1) normal, (2) infiltrates other than endpoint pneumonia, and (3) lobar consolidation or pleural effusion (endpoint pneumonia). PCR = polymerase chain reaction; WBC = white blood cells; Culture = bacterial blood or pleural fluid culture for respiratory pathogens (contaminants excluded).

**Fig. S2 Heatmaps of unsupervised hierarchical clustering of all 1107 proteins and BvVM markers (n=151)**

**(A)** Heatmap from unsupervised hierarchical clustering of all 1107 proteins measured in the space of all 171 single etiology samples from the training and validation set.

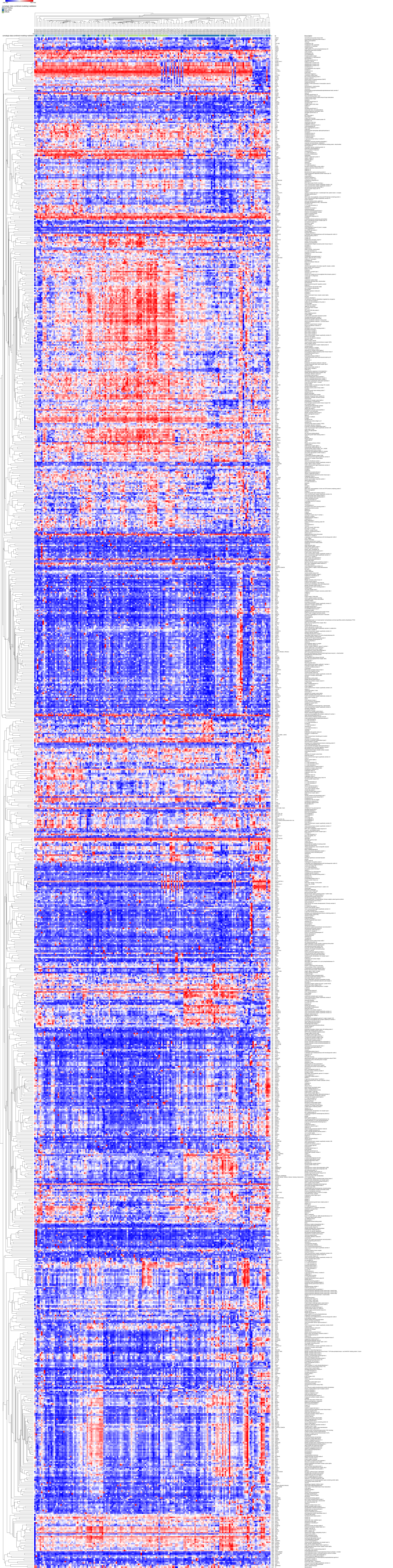
row min
row max
somalogic-data-combined-modeling+validation
BACTERIA
MALARIA
VIRUS
id
somalogic-data-combined-modeling+validation

(**B**) Heatmap for the Bacteria vs Virus or Malaria model with all 151 significant markers, hierarchically clustered with all single etiology bacteria, virus, and control samples from the training and validation set (n=171).

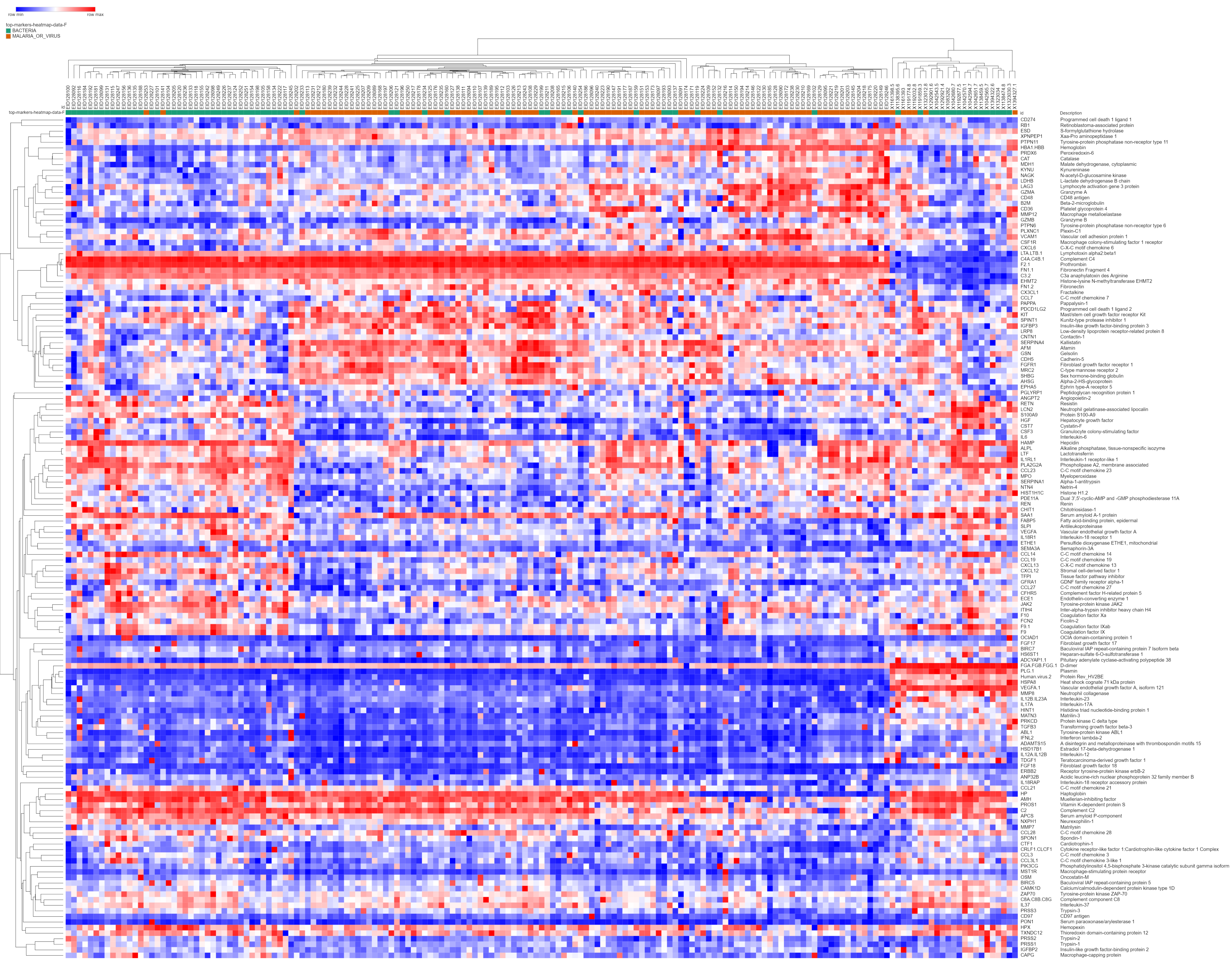
row min
row max
top-markers-heatmap-data-F
BACTERIA
MALARIA_OR_VIRUS
id
top-markers-heatmap-data-F
id
Description
CD274 Programmed cell death 1 ligand 1
RB1 Retinoblastoma-associated protein
ESD S-formylglutathione hydrolase
XPNPEP1 Xaa-Pro aminopeptidase 1
PTPN11 Tyrosine-protein phosphatase non-receptor type 11
HBA1.HBB Hemoglobin
PRDX6 Peroxiredoxin-6
CAT Catalase
MDH1 Malate dehydrogenase, cytoplasmic
KYNU Kynureninase
NAGK N-acetyl-D-glucosamine kinase
LDHB L-lactate dehydrogenase B chain
LAG3 Lymphocyte activation gene 3 protein
GZMA Granzyme A
CD48 CD48 antigen
B2M Beta-2-microglobulin
CD36 Platelet glycoprotein 4
MMP12 Macrophage metalloelastase
GZMB Granzyme B
PTPN6 Tyrosine-protein phosphatase non-receptor type 6
PLXNC1 Plexin-C1
VCAM1 Vascular cell adhesion protein 1
CSF1R Macrophage colony-stimulating factor 1 receptor
CXCL6 C-X-C motif chemokine 6
LTA.LTB.1 Lymphotoxin alpha2:beta1
C4A.C4B.1 Complement C4
F2.1 Prothrombin
FN1.1 Fibronectin Fragment 4
C3.2 C3a anaphylatoxin des Arginine
EHMT2 Histone-lysine N-methyltransferase EHMT2
FN1.2 Fibronectin
CX3CL1 Fractalkine
CCL7 C-C motif chemokine 7
PAPPA Pappalysin-1
PDCD1LG2 Programmed cell death 1 ligand 2
KIT Mast/stem cell growth factor receptor Kit
SPINT1 Kunitz-type protease inhibitor 1
IGFBP3 Insulin-like growth factor-binding protein 3
LRP8 Low-density lipoprotein receptor-related protein 8
CNTN1 Contactin-1
SERPINA4 Kallistatin
AFM Afamin
GSN Gelsolin
CDH5 Cadherin-5
FGFR1 Fibroblast growth factor receptor 1
MRC2 C-type mannose receptor 2
SHBG Sex hormone-binding globulin
AHSG Alpha-2-HS-glycoprotein
EPHA5 Ephrin type-A receptor 5
PGLYRP1 Peptidoglycan recognition protein 1
ANGPT2 Angiopoietin-2
RETN Resistin
LCN2 Neutrophil gelatinase-associated lipocalin
S100A9 Protein S100-A9
HGF Hepatocyte growth factor
CST7 Cystatin-F
CSF3 Granulocyte colony-stimulating factor
IL6 Interleukin-6
HAMP Hepcidin
ALPL Alkaline phosphatase, tissue-nonspecific isozyme
LTF Lactotransferrin
IL1RL1 Interleukin-1 receptor-like 1
PLA2G2A Phospholipase A2, membrane associated
CCL23 C-C motif chemokine 23
MPO Myeloperoxidase
SERPINA1 Alpha-1-antitrypsin
NTN4 Netrin-4
HIST1H1C Histone H1.2
PDE11A Dual 3',5'-cyclic-AMP and -GMP phosphodiesterase 11A
REN Renin
CHIT1 Chitotriosidase-1
SAA1 Serum amyloid A-1 protein
FABP5 Fatty acid-binding protein, epidermal
SLPI Antileukoproteinase
VEGFA Vascular endothelial growth factor A
IL18R1 Interleukin-18 receptor 1
ETHE1 Persulfide dioxygenase ETHE1, mitochondrial
SEMA3A Semaphorin-3A
CCL14 C-C motif chemokine 14
CCL19 C-C motif chemokine 19
CXCL13 C-X-C motif chemokine 13
CXCL12 Stromal cell-derived factor 1
TFPI Tissue factor pathway inhibitor
GFRA1 GDNF family receptor alpha-1
CCL27 C-C motif chemokine 27
CFHR5 Complement factor H-related protein 5
ECE1 Endothelin-converting enzyme 1
JAK2 Tyrosine-protein kinase JAK2
ITIH4 Inter-alpha-trypsin inhibitor heavy chain H4
F10 Coagulation factor Xa
FCN2 Ficolin-2
F9.1 Coagulation factor IXab
F9 Coagulation factor IX
OCIAD1 OCIA domain-containing protein 1
FGF17 Fibroblast growth factor 17
BIRC7 Baculoviral IAP repeat-containing protein 7 Isoform beta
HS6ST1 Heparan-sulfate 6-O-sulfotransferase 1
ADCYAP1.1 Pituitary adenylate cyclase-activating polypeptide 38
FGA.FGB.FGG.1 D-dimer
PLG.1 Plasmin
Human.virus.2 Protein Rev_HV2BE
HSPA8 Heat shock cognate 71 kDa protein
VEGFA.1 Vascular endothelial growth factor A, isoform 121
MMP8 Neutrophil collagenase
IL12B.IL23A Interleukin-23
IL17A Interleukin-17A
HINT1 Histidine triad nucleotide-binding protein 1
MATN3 Matrilin-3
PRKCD Protein kinase C delta type
TGFB3 Transforming growth factor beta-3
ABL1 Tyrosine-protein kinase ABL1
IFNL2 Interferon lambda-2
ADAMTS15 A disintegrin and metalloproteinase with thrombospondin motifs 15
HSD17B1 Estradiol 17-beta-dehydrogenase 1
IL12A.IL12B Interleukin-12
TDGF1 Teratocarcinoma-derived growth factor 1
FGF18 Fibroblast growth factor 18
ERBB2 Receptor tyrosine-protein kinase erbB-2
ANP32B Acidic leucine-rich nuclear phosphoprotein 32 family member B
IL18RAP Interleukin-18 receptor accessory protein
CCL21 C-C motif chemokine 21
HP Haptoglobin
AMH Muellerian-inhibiting factor
PROS1 Vitamin K-dependent protein S
C2 Complement C2
APCS Serum amyloid P-component
NXPH1 Neurexophilin-1
MMP7 Matrilysin
CCL28 C-C motif chemokine 28
SPON1 Spondin-1
CTF1 Cardiotrophin-1
CRLF1.CLCF1 Cytokine receptor-like factor 1:Cardiotrophin-like cytokine factor 1 Complex
CCL3 C-C motif chemokine 3
CCL3L1 C-C motif chemokine 3-like 1
PIK3CG Phosphatidylinositol 4,5-bisphosphate 3-kinase catalytic subunit gamma isoform
MST1R Macrophage-stimulating protein receptor
OSM Oncostatin-M
BIRC5 Baculoviral IAP repeat-containing protein 5
CAMK1D Calcium/calmodulin-dependent protein kinase type 1D
ZAP70 Tyrosine-protein kinase ZAP-70
C8A.C8B.C8G Complement component C8
IL37 Interleukin-37
PRSS3 Trypsin-3
CD97 CD97 antigen
PON1 Serum paraoxonase/arylesterase 1
HPX Hemopexin
TXNDC12 Thioredoxin domain-containing protein 12
PRSS2 Trypsin-2
PRSS1 Trypsin-1
IGFBP2 Insulin-like growth factor-binding protein 2
CAPG Macrophage-capping protein

**Fig. S3 Protein marker overlap of the 30 highest ranking markers.**

SOMAScan Bacteria vs Virus (BvV) and Bacteria vs Virus and Malaria (BvVM) marker sets.

| Comparison | Markers | Venn Diagram |
|---|---|---|
| SOMAScan (BvV) and SOMAScan (BvVM) | CSF3<br>CTF1<br>CX3CL1<br>CXCL13<br>ESD<br>FABP5<br>HGF<br>IL1RL1<br>IL6<br>LCN2<br>MPO<br>PIK3CG<br>PLA2G2A<br>RETN<br>S100A9<br>SERPINA4<br>SLPI | 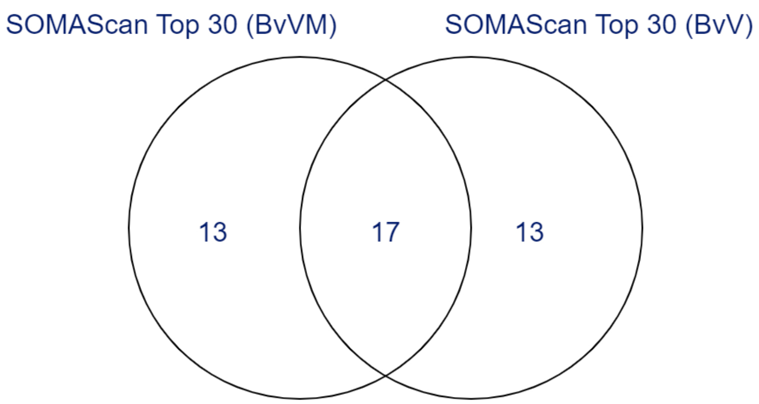 |

**Fig. S4 Box and whisker plots of the 100 Top markers.**

Distribution of the logarithm of Relative Fluorescence Units (RFU) for BvV and BvVM marker sets. RFUs are directly proportional to the amount of target protein in the initial sample. B = Bacteria, V = Virus, V+M = Virus or Malaria

**(A) Box and** whisker plots of 100 Top Markers in Bacteria vs Virus or Malaria

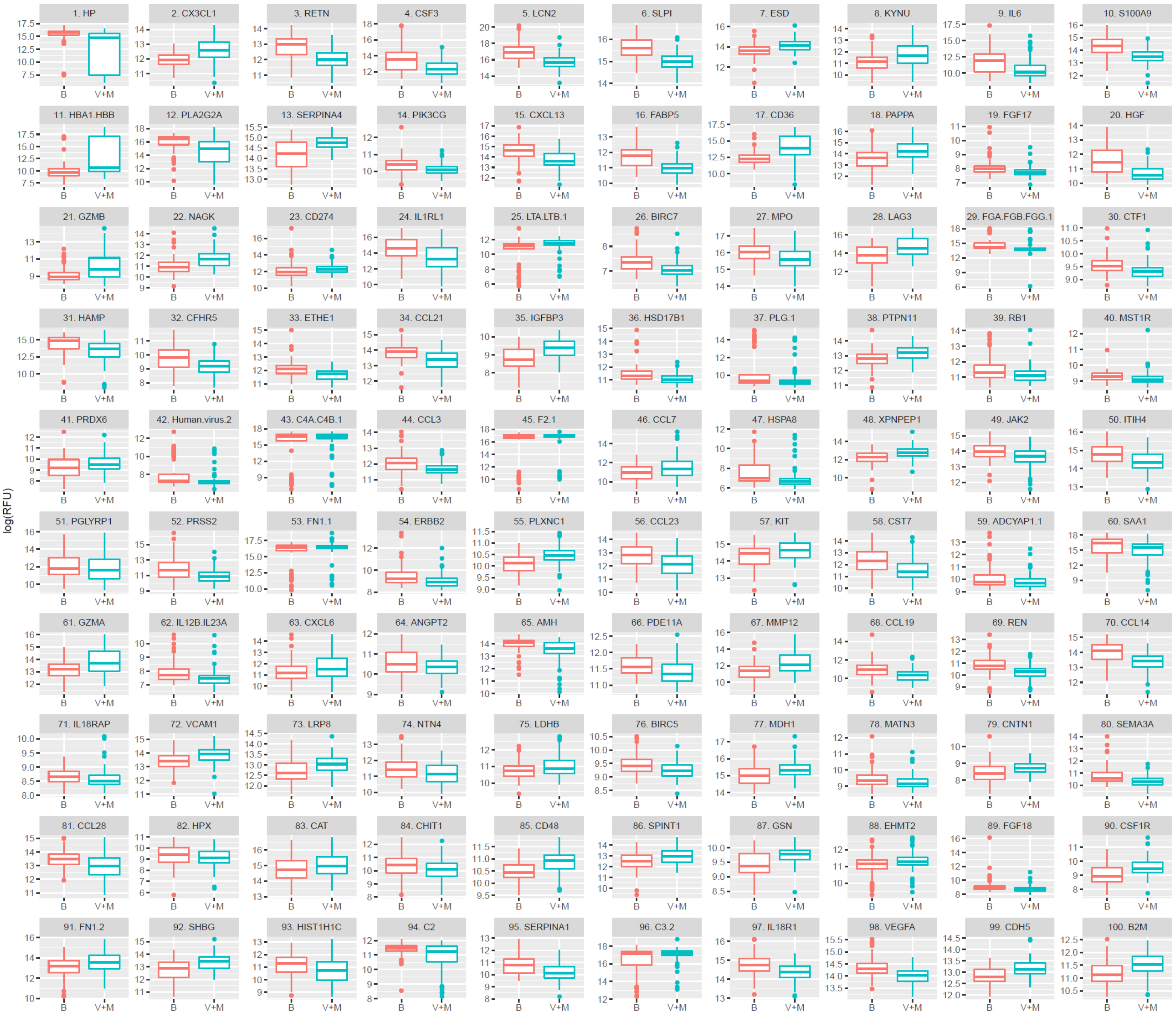

**(B)** Box and whisker plots of 100 Top Markers in Bacteria vs Virus

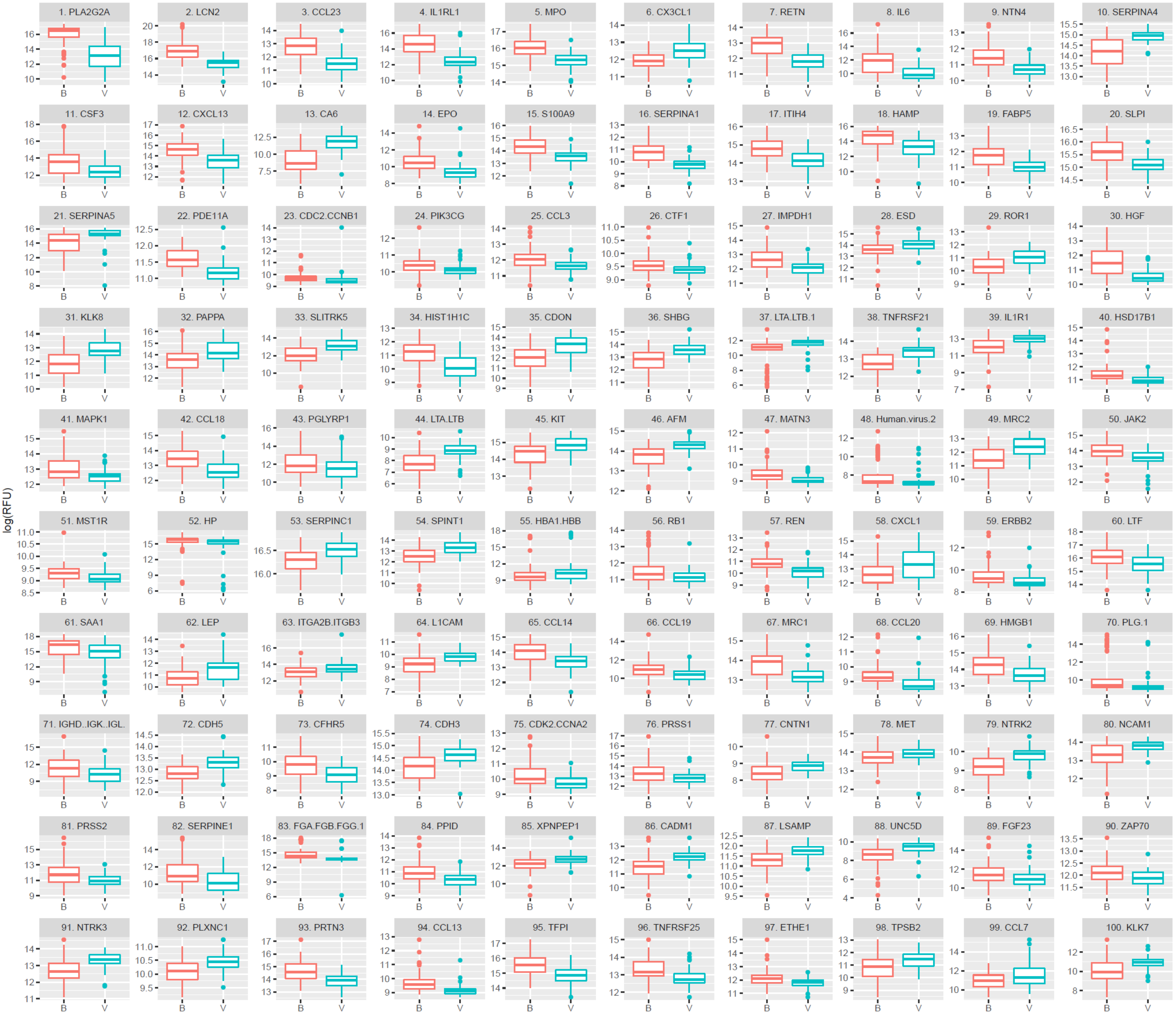

**Fig. S5 Model accuracy when switching out surrogates for comparisons of interests**

Model classification accuracy assessed on validation data for Bacteria vs Virus and Bacteria vs Virus or Malaria using two different classification algorithms (Random Forest and Elastic Net). The plots show classifier accuracy with increasing numbers of biomarkers used in training the model. The panels show how model performance changes when 10% or 20% of markers are substituted with their corresponding surrogate markers.

**(A) Bacteria vs Virus** with 5, 10, 15, 25, 50, 100 and all (219) markers

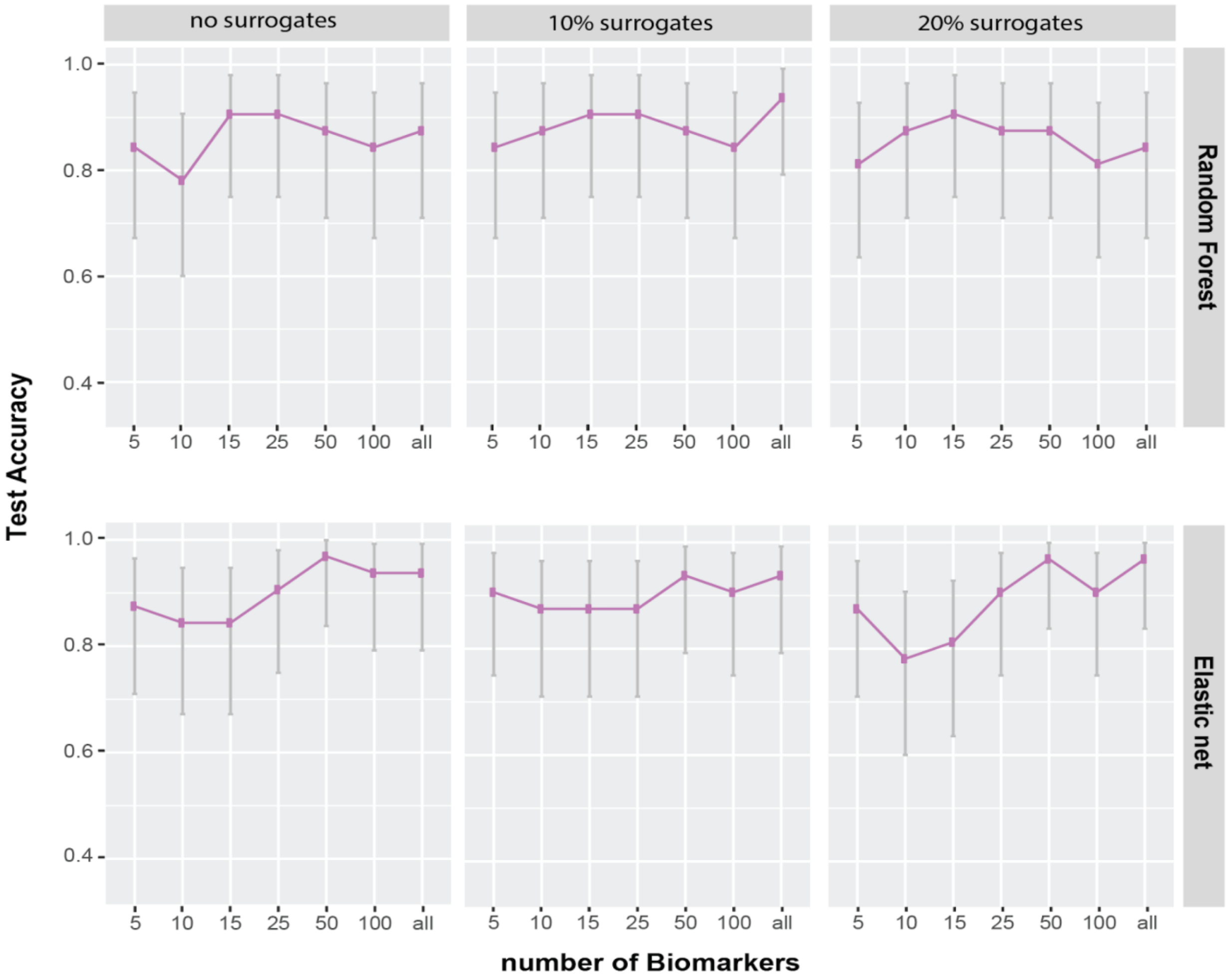

**(B) Bacteria vs Malaria or Virus** with 5, 10, 15, 25, 50, 100 and all (151) markers

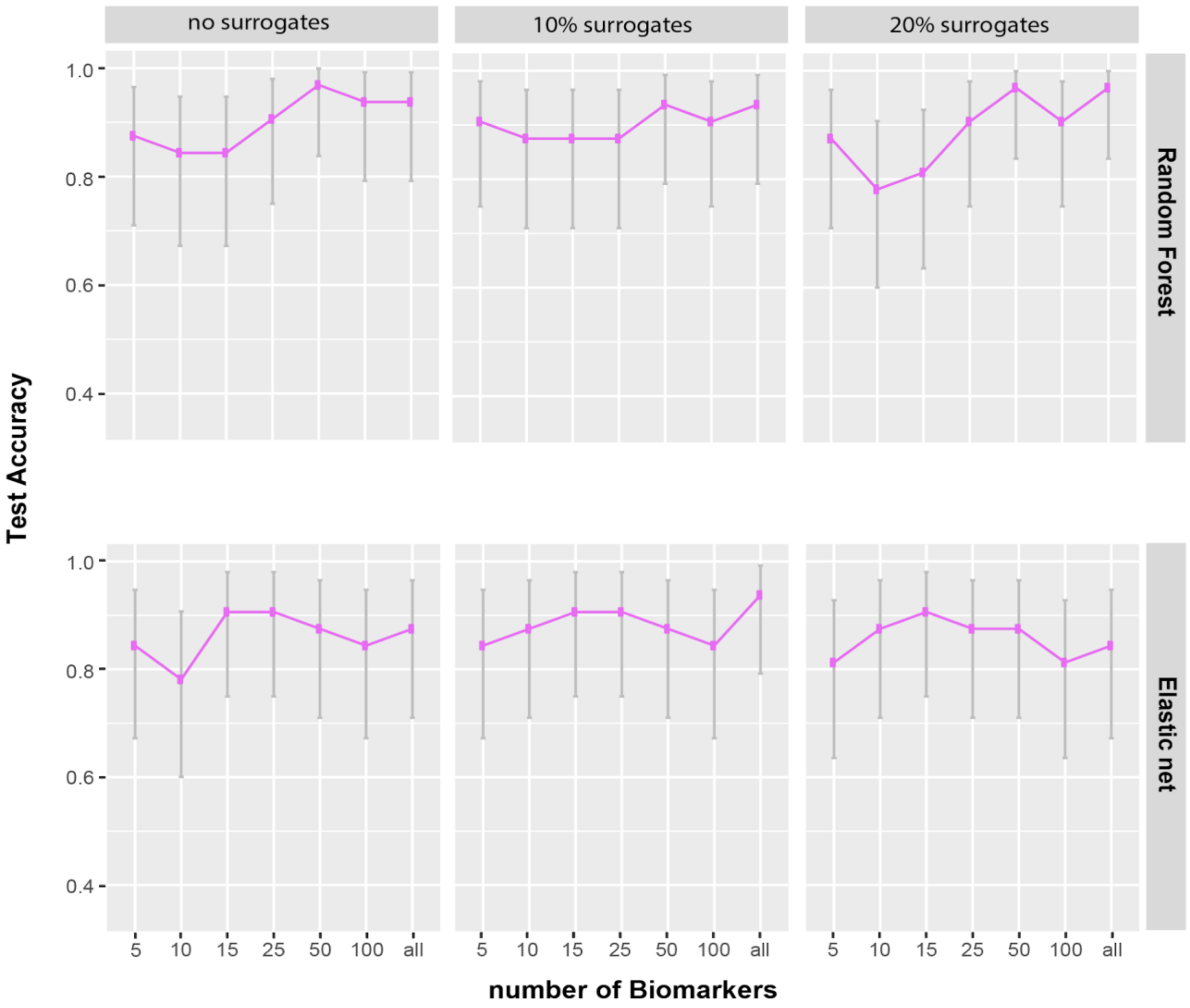

**Fig. S6 RBM multiplexed Immunoassay, RNA-sequencing and SOMAscan assay comparisons with Venn diagrams and marker lists**

(**A**) Table of differential markers (genes) shared between this study and the previous RNA-Sequencing study of the same population.[1,2] SOMAScan protein markers were converted into their gene names to compare markers between studies.

Marker sets (genes) represented are SOMAScan BvV (n=226), SOMAScan BvVM (n=156), and RNA-Sequencing BvVM, Malaria vs Virus or Malaria (MvBV), and Virus vs Bacteria or Malaria (VvBM) (n=431, n=99, and n=70, respectively).

| **Comparison** | **Markers** | **Venn Diagram** |
|---|---|---|
| RNA-Sequencing (BvVM, MvVB, VvBM) and SOMAScan (BvVM)<br><br>The 25 gene marker overlap translates to 24 protein markers<br><br>[+]HBA1 and HBB is hemoglobin<br><br>*C3a in SOMAScan | ALPL<br>C3*<br>CCL3<br>CD274<br>CHIT1<br>CST7<br>GZMB<br>HBA1[+]<br>HBB[+]<br>HGF<br>HP<br>IL18R1<br>IL18RAP<br>LAG3<br>LCN2<br>LTF<br>MMP8<br>MPO<br>OSM<br>PGLYRP1<br>PLXNC1 | 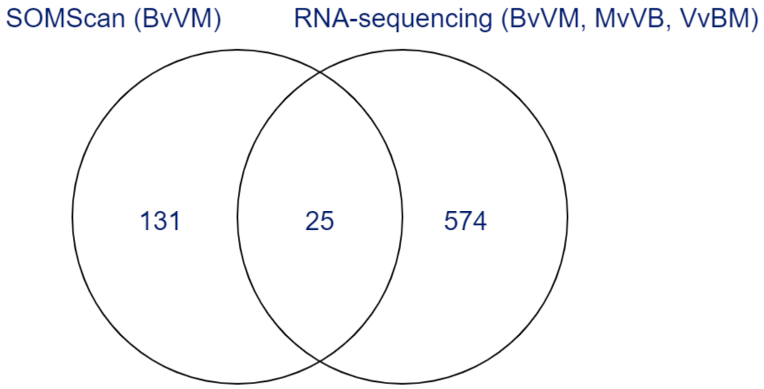 |

| | | |
|---|---|---|
| | RETN<br>S100A9<br>SERPINA1<br>SLPI | |
| SOMAScan (BvVM) and RNA-Sequencing (BvVM)<br><br>*depicted in Fig. 4 B* | ALPL<br>CHIT1<br>CST7<br>HGF<br>HP<br>IL18R1<br>IL18RAP<br>LCN2<br>LTF<br>MMP8<br>MPO<br>OSM<br>PGLYRP1<br>PLXNC1<br>RETN<br>S100A9<br>SERPINA1<br>SLPI | 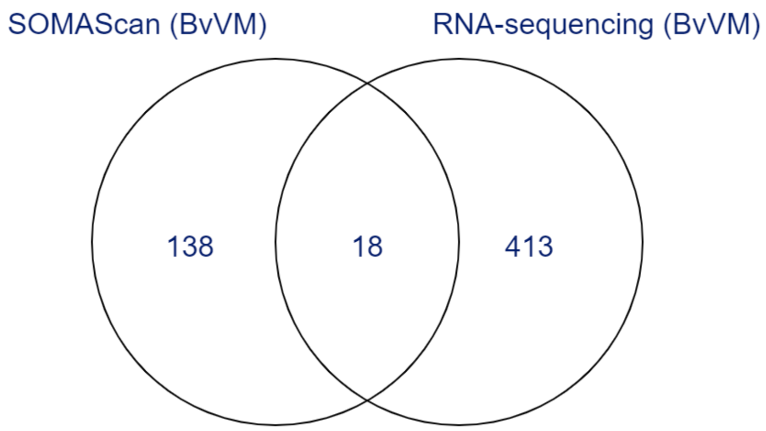 |

| | | |
|---|---|---|
| RNA-Sequencing (BvVM) and SOMAScan (BvV) | ALPL<br>BPI<br>CHIT1<br>CST7<br>HGF<br>HP<br>IL18R1<br>IL1R1<br>ITGA2B<br>ITGB3<br>LCN2<br>LTF<br>MAPK14<br>MPO<br>PGD<br>PGLYRP1<br>PLXNC1<br>RETN<br>S100A9<br>SERPINA1<br>SLPI<br>SMPDL3A | 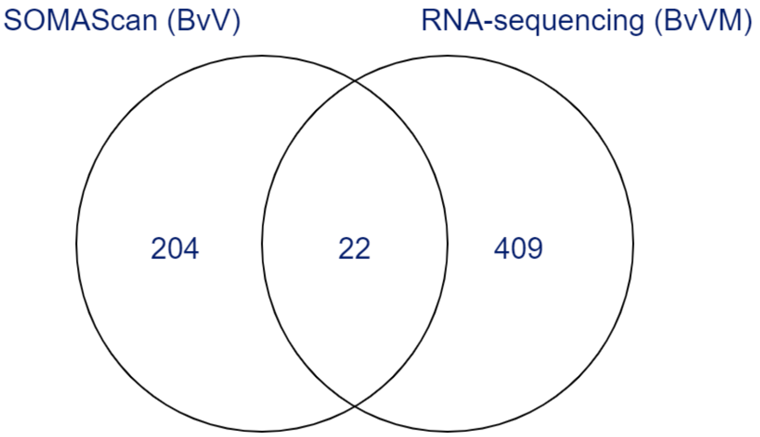 |
| RNA-Sequencing (MvVB) and SOMAScan (BvVM)<br><br>7 gene marker overlap translates to 6 protein markers<br><br>*depicted in Supplemental File 6C*<br><br>*C3a in SOMAScan<br>$^{+}$ HBA1 and HBB is hemoglobin | C3*<br>CCL3<br>CD274<br>GZMB<br>HBA1$^{+}$<br>HBB$^{+}$<br>LAG3 | 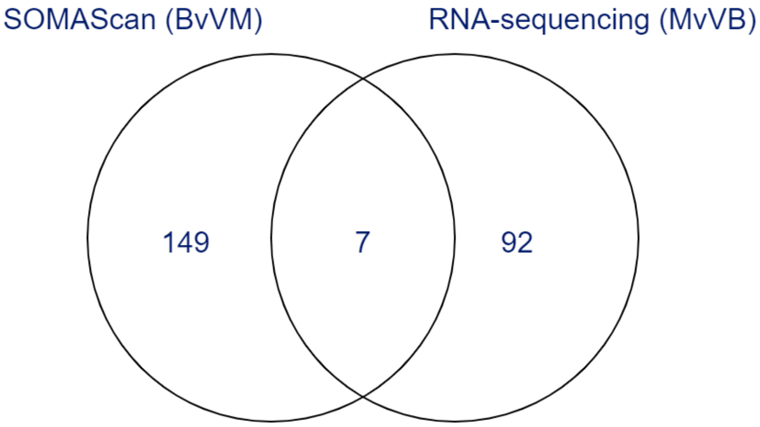 |

(**B**) Table of differential markers (genes) shared between this study and previous RBM protein multiplex immunoassay of the same population [1,2]. RBM and SOMAScan protein aliases were converted into their gene names to compare markers between studies.

Marker sets (genes) represented are SOMAScan BvV (n=226), SOMAScan BvVM (n=156), RBM multiplex assay 2-class comparisons (BvV, BvM, VvM with n=15, n=9, and n=22, respectively) and RBM multiplex assay 3-class comparison Bacteria vs Virus vs Malaria (n=16).

| **Comparison** | **Markers** | **Venn Diagram** |
|---|---|---|
| SOMAScan (BvV) and RBM (BvV)<br><br>*depicted in Fig. 3* | CKM<br>HP<br>IL6<br>MPO<br>SERPINA1 | 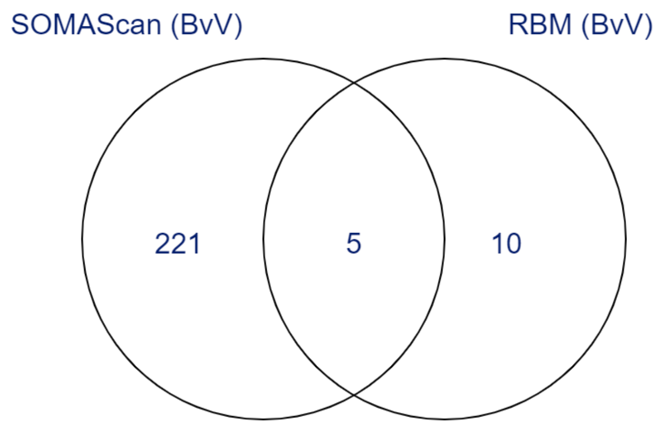 |
| SOMAScan (BvVM) and RBM (BvV, BvM) | APCS<br>HP<br>IL6<br>MPO<br>SERPINA1<br>VCAM1 | 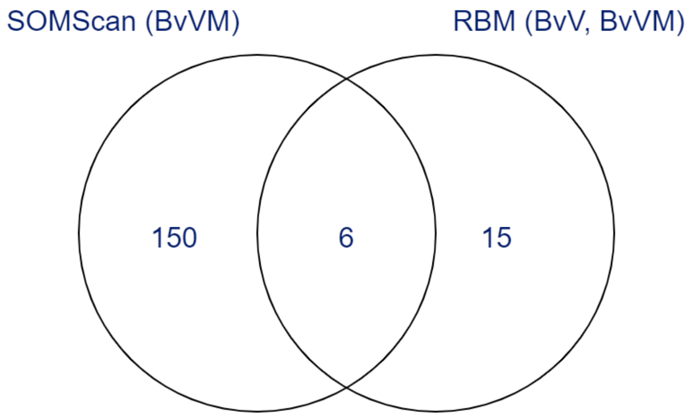 |

**Fig. S7 Heatmap of all 24 overlapping SOMAScan (BvVM) and RNA-sequencing markers.** Hierarchically clustered heatmap of normalized SOMAscan expression values for the 24 overlapping protein markers from the RNA-Sequencing study and the SOMAScan Bacteria vs Malaria or Virus comparison in the space of all available single etiology samples used in this study (n=167). The markers depicted are ALPL, CHIT1, C3a, CCL3, CD274, CST7, GZMB, HBA1.HBB (hemoglobin), HGF, HP, IL18R1, IL18RAP, LAG3, LCN2, LTF, MMP8, MPO, OSM, PGLYRP1, PLXNC1, RETN, S100A9, SERPINA1, and SLPI.

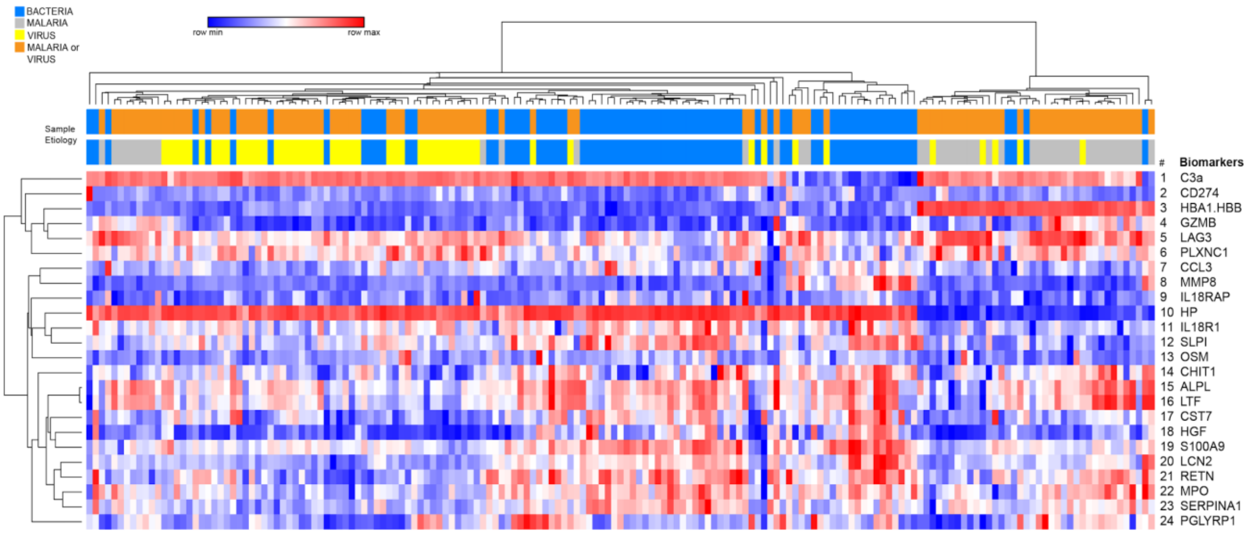

**Fig. S8 Malarial markers shared between SOMAScan and RNA-sequencing.**

Differential markers shared between this study and previous RNA-sequencing study of the same population. Shown are six protein markers shared between this study (SOMAScan BvVM) and RNA-sequencing Malaria vs Bacteria or Virus (MvVB). Red = upregulated in bacterial samples, black = upregulated in malarial samples, dark blue = upregulated in malaria or virus samples.

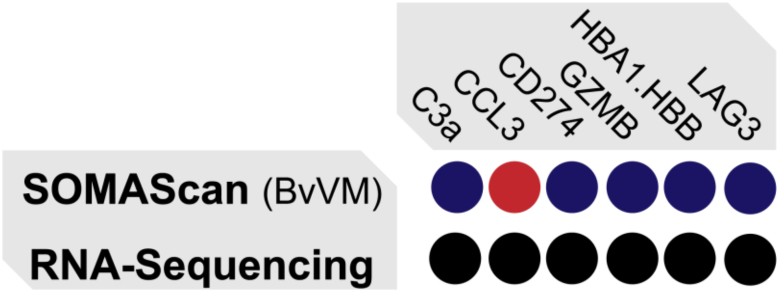

**Fig. S9 Enrichment Heatmap of the three-way comparison (Bacteria vs Malaria vs Virus).**

In this comparison individual protein markers could be shared by two, but not all three etiologies.

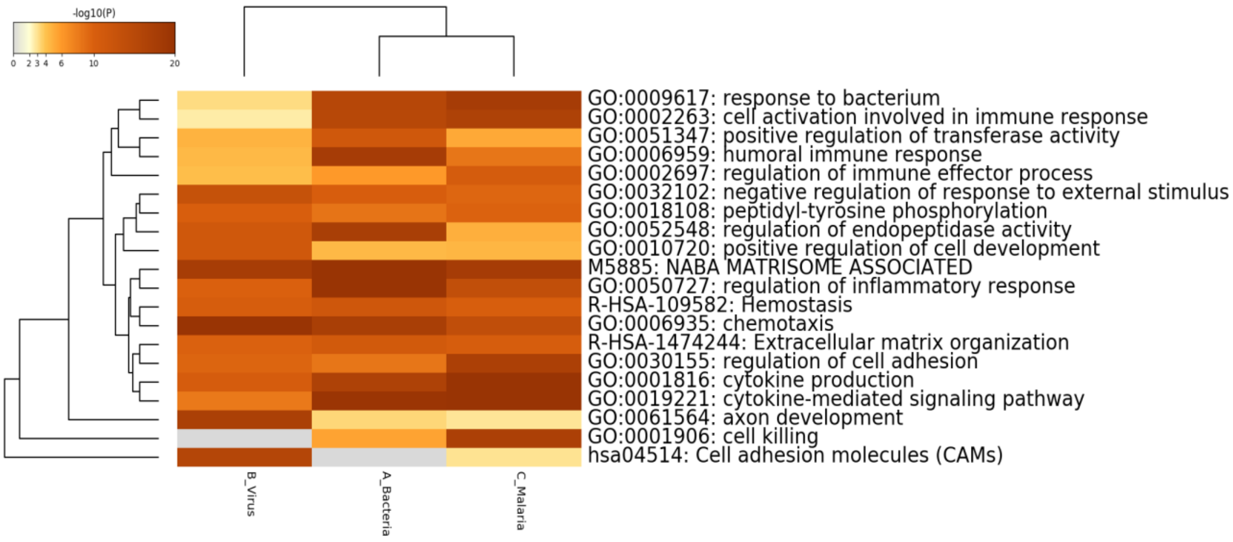

**Fig. S10 Heatmap of the Bacteria vs Virus model.** (A) Hierarchically clustered heatmap of normalized SOMAscan expression values for 219 significant markers (FDR<0.01) from the SOMAScan Bacteria vs Virus (BvV) comparison in the space of all single etiology bacterial and viral samples in this study.

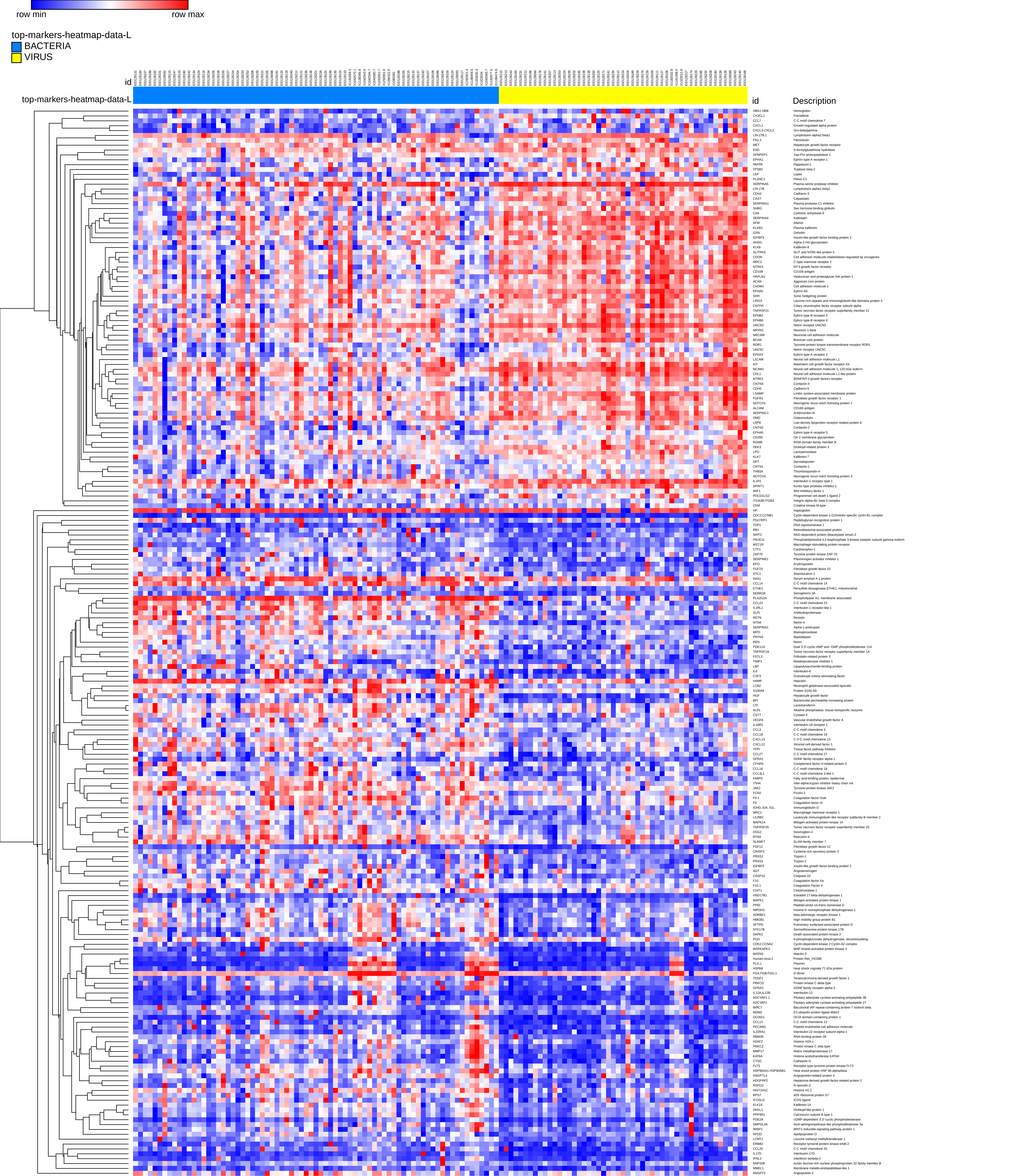
row min
row max
top-markers-heatmap-data-L
BACTERIA
VIRUS
id
top-markers-heatmap-data-L
id
Description
HBA1.HBB Hemoglobin
CX3CL1 Fractalkine
CCL7 C-C motif chemokine 7
CXCL1 Growth-regulated alpha protein
CXCL3.CXCL2 Gro-beta/gamma
LTA.LTB.1 Lymphotoxin alpha2:beta1
FN1.2 Fibronectin
MET Hepatocyte growth factor receptor
ESD S-formylglutathione hydrolase
XPNPEP1 Xaa-Pro aminopeptidase 1
EPHA1 Ephrin type-A receptor 1
PAPPA Pappalysin-1
TPSB2 Tryptase beta-2
LEP Leptin
PLXNC1 Plexin-C1
SERPINA5 Plasma serine protease inhibitor
LTA.LTB Lymphotoxin alpha1:beta2
CDH3 Cadherin-3
CAST Calpastatin
SERPING1 Plasma protease C1 inhibitor
SHBG Sex hormone-binding globulin
CA6 Carbonic anhydrase 6
SERPINA4 Kallistatin
AFM Afamin
KLKB1 Plasma kallikrein
GSN Gelsolin
IGFBP3 Insulin-like growth factor-binding protein 3
AHSG Alpha-2-HS-glycoprotein
KLK8 Kallikrein-8
SLITRK5 SLIT and NTRK-like protein 5
CDON Cell adhesion molecule-related/down-regulated by oncogenes
MRC2 C-type mannose receptor 2
NTRK3 NT-3 growth factor receptor
CD109 CD109 antigen
HAPLN1 Hyaluronan and proteoglycan link protein 1
ACAN Aggrecan core protein
CADM1 Cell adhesion molecule 1
EFNA5 Ephrin-A5
SHH Sonic hedgehog protein
LRIG3 Leucine-rich repeats and immunoglobulin-like domains protein 3
CNTFR Ciliary neurotrophic factor receptor subunit alpha
TNFRSF21 Tumor necrosis factor receptor superfamily member 21
EPHB2 Ephrin type-B receptor 2
EPHB6 Ephrin type-B receptor 6
UNC5D Netrin receptor UNC5D
NRXN1 Neurexin-1-beta
NRCAM Neuronal cell adhesion molecule
BCAN Brevican core protein
ROR1 Tyrosine-protein kinase transmembrane receptor ROR1
UNC5C Netrin receptor UNC5C
EPHA2 Ephrin type-A receptor 2
L1CAM Neural cell adhesion molecule L1
KIT Mast/stem cell growth factor receptor Kit
NCAM1 Neural cell adhesion molecule 1, 120 kDa isoform
CHL1 Neural cell adhesion molecule L1-like protein
NTRK2 BDNF/NT-3 growth factors receptor
CNTN4 Contactin-4
CDH5 Cadherin-5
LSAMP Limbic system-associated membrane protein
FGFR1 Fibroblast growth factor receptor 1
NOTCH1 Neurogenic locus notch homolog protein 1
ALCAM CD166 antigen
SERPINC1 Antithrombin-III
OMD Osteomodulin
LRP8 Low-density lipoprotein receptor-related protein 8
CNTN2 Contactin-2
EPHA5 Ephrin type-A receptor 5
CD200 OX-2 membrane glycoprotein
RGMB RGM domain family member B
DKK3 Dickkopf-related protein 3
LPO Lactoperoxidase
KLK7 Kallikrein-7
DPT Dermatopontin
CNTN1 Contactin-1
THBS4 Thrombospondin-4
NOTCH3 Neurogenic locus notch homolog protein 3
IL1R1 Interleukin-1 receptor type 1
SPINT1 Kunitz-type protease inhibitor 1
WIF1 Wnt inhibitory factor 1
PDCD1LG2 Programmed cell death 1 ligand 2
ITGA2B.ITGB3 Integrin alpha-IIb: beta-3 complex
CKM Creatine kinase M-type
HP Haptoglobin
CDC2.CCNB1 Cyclin-dependent kinase 1:G2/mitotic-specific cyclin-B1 complex
PGLYRP1 Peptidoglycan recognition protein 1
TOP1 DNA topoisomerase 1
RB1 Retinoblastoma-associated protein
SIRT2 NAD-dependent protein deacetylase sirtuin-2
PIK3CG Phosphatidylinositol 4,5-bisphosphate 3-kinase catalytic subunit gamma isoform
MST1R Macrophage-stimulating protein receptor
CTF1 Cardiotrophin-1
ZAP70 Tyrosine-protein kinase ZAP-70
SERPINE1 Plasminogen activator inhibitor 1
EPO Erythropoietin
FGF23 Fibroblast growth factor 23
STC1 Stanniocalcin-1
SAA1 Serum amyloid A-1 protein
CCL14 C-C motif chemokine 14
ETHE1 Persulfide dioxygenase ETHE1, mitochondrial
SEMA3A Semaphorin-3A
PLA2G2A Phospholipase A2, membrane associated
CCL23 C-C motif chemokine 23
IL1RL1 Interleukin-1 receptor-like 1
SLPI Antileukoproteinase
RETN Resistin
NTN4 Netrin-4
SERPINA1 Alpha-1-antitrypsin
MPO Myeloperoxidase
PRTN3 Myeloblastin
REN Renin
PDE11A Dual 3',5'-cyclic-AMP and -GMP phosphodiesterase 11A
TNFRSF1A Tumor necrosis factor receptor superfamily member 1A
FSTL3 Follistatin-related protein 3
TIMP1 Metalloproteinase inhibitor 1
LBP Lipopolysaccharide-binding protein
IL6 Interleukin-6
CSF3 Granulocyte colony-stimulating factor
HAMP Hepcidin
LCN2 Neutrophil gelatinase-associated lipocalin
S100A9 Protein S100-A9
HGF Hepatocyte growth factor
BPI Bactericidal permeability-increasing protein
LTF Lactotransferrin
ALPL Alkaline phosphatase, tissue-nonspecific isozyme
CST7 Cystatin-F
VEGFA Vascular endothelial growth factor A
IL18R1 Interleukin-18 receptor 1
CCL3 C-C motif chemokine 3
CCL19 C-C motif chemokine 19
CXCL13 C-X-C motif chemokine 13
CXCL12 Stromal cell-derived factor 1
TFPI Tissue factor pathway inhibitor
CCL27 C-C motif chemokine 27
GFRA1 GDNF family receptor alpha-1
CFHR5 Complement factor H-related protein 5
CCL18 C-C motif chemokine 18
CCL3L1 C-C motif chemokine 3-like 1
FABP5 Fatty acid-binding protein, epidermal
ITIH4 Inter-alpha-trypsin inhibitor heavy chain H4
JAK2 Tyrosine-protein kinase JAK2
FCN2 Ficolin-2
F9.1 Coagulation factor IXab
F9 Coagulation factor IX
IGHD..IGK..IGL. Immunoglobulin D
MRC1 Macrophage mannose receptor 1
LILRB2 Leukocyte immunoglobulin-like receptor subfamily B member 2
MAPK14 Mitogen-activated protein kinase 14
TNFRSF25 Tumor necrosis factor receptor superfamily member 25
DSG2 Desmoglein-2
RTN4 Reticulon-4
SLAMF7 SLAM family member 7
FGF12 Fibroblast growth factor 12
CRISP3 Cysteine-rich secretory protein 3
PRSS1 Trypsin-1
PRSS2 Trypsin-2
IGFBP2 Insulin-like growth factor-binding protein 2
AGT Angiotensinogen
CASP10 Caspase-10
F10 Coagulation factor Xa
F10.1 Coagulation Factor X
CHIT1 Chitotriosidase-1
HSD17B1 Estradiol 17-beta-dehydrogenase 1
MAPK1 Mitogen-activated protein kinase 1
PPID Peptidyl-prolyl cis-trans isomerase D
IMPDH1 Inosine-5'-monophosphate dehydrogenase 1
ADRBK1 beta-adrenergic receptor kinase 1
HMGB1 High mobility group protein B1
SFTPD Pulmonary surfactant-associated protein D
STK17B Serine/threonine-protein kinase 17B
DAPK2 Death-associated protein kinase 2
PGD 6-phosphogluconate dehydrogenase, decarboxylating
CDK2.CCNA2 Cyclin-dependent kinase 2:Cyclin-A2 complex
MAPKAPK3 MAP kinase-activated protein kinase 3
MATN3 Matrilin-3
Human.virus.2 Protein Rev_HV2BE
PLG.1 Plasmin
HSPA8 Heat shock cognate 71 kDa protein
FGA.FGB.FGG.1 D-dimer
TDGF1 Teratocarcinoma-derived growth factor 1
PRKCD Protein kinase C delta type
GFRA3 GDNF family receptor alpha-3
IL12A.IL12B Interleukin-12
ADCYAP1.1 Pituitary adenylate cyclase-activating polypeptide 38
ADCYAP1 Pituitary adenylate cyclase-activating polypeptide 27
BIRC7 Baculoviral IAP repeat-containing protein 7 Isoform beta
MDM2 E3 ubiquitin-protein ligase Mdm2
OCIAD1 OCIA domain-containing protein 1
CCL13 C-C motif chemokine 13
PECAM1 Platelet endothelial cell adhesion molecule
IL22RA1 Interleukin-22 receptor subunit alpha-1
RBM39 RNA-binding protein 39
H2AFZ Histone H2A.z
PRKCZ Protein kinase C zeta type
MMP17 Matrix metalloproteinase-17
KAT6A Histone acetyltransferase KAT6A
CTSG Cathepsin G
FLT3 Receptor-type tyrosine-protein kinase FLT3
HSP90AA1.HSP90AB1 Heat shock protein HSP 90-alpha/beta
ANGPTL4 Angiopoietin-related protein 4
HDGFRP2 Hepatoma-derived growth factor-related protein 2
RSPO2 R-spondin-2
HIST1H1C Histone H1.2
RPS7 40S ribosomal protein S7
ICOSLG ICOS ligand
KLK14 Kallikrein-14
DKKL1 Dickkopf-like protein 1
PPP3R1 Calcineurin subunit B type 1
PDE2A cGMP-dependent 3',5'-cyclic phosphodiesterase
SMPDL3A Acid sphingomyelinase-like phosphodiesterase 3a
WISP1 WNT1-inducible-signaling pathway protein 1
APOD Apolipoprotein D
LCMT1 Leucine carboxyl methyltransferase 1
ERBB2 Receptor tyrosine-protein kinase erbB-2
CCL20 C-C motif chemokine 20
IL17D Interleukin-17D
IFNL2 Interferon lambda-2
ANP32B Acidic leucine-rich nuclear phosphoprotein 32 family member B
MMEL1 Membrane metallo-endopeptidase-like 1
ANGPT2 Angiopoietin-2

#### Tables

**Table S1 Patient demographic and disease characteristics at admission for Bacterial and PCR Bacterial samples.**

The “Bacteria” group includes blood or pleural fluid culture positive samples and “probable bacteria” samples that were culture negative but had leukocytosis and radiographic endpoint pneumonia and were independently judged by two clinical experts as probable bacterial infections.

**Patient Demographic and Disease Characteristics at admission**

| Features on admission (signs, symptoms, and laboratory results) | Bacteria | No. | PCR Bacteria$^{\Omega}$ | No. | P$^{\dagger}$ |
|---|---|---|---|---|---|
| Age (month), mean (SD) | 30.5 (30.9) | 42 | 28.4 (27.5) | 27 | 0.87 |
| Female sex, n (%) | 19 (45.2) | 42 | 12 (44.4) | 27 | 0.79 |
| **Clinical Examination results on arrival** | | | | | |
| Weight (kg), mean (SD) | 10.1 (4.6) | 42 | 10.2 (5.2) | 27 | 0.69 |
| Height (cm), mean (SD) | 81.3 (18.8) | 42 | 78.8 (18.8) | 26 | 0.57 |
| MUAC (cm), mean (SD) | 13.4 (2.1) | 42 | 13.56 (1.8) | 27 | 0.86 |
| Temperature (°C), mean (SD) | 38.5 (1.0) | 42 | 37.69 (1.3) | 27 | 0.050 |
| Respiratory rate (cycles per min), mean (SD) | 61 (14.9) | 42 | 60 (14.7) | 27 | 0.77 |
| **Nutritional status** | | | | | |
| WAZ > -1 SD | 11 (26.2) | 42 | 6 (22.2) | 27 | |
| WAZ -1 SD to -3 SD, n (%) (low to severe underweight) | 20 (47.6) | 42 | 16 (59.3) | 27 | |
| WAZ < -3 SD, n (%) (severe underweight) | 11 (26.2) | 42 | 5 (18.5) | 27 | |
| WAZ: Mean (SD) | -2 (1.8) | 42 | -2 (1.7) | 27 | 0.92 |
| **Anaemia status on admission** | | | | | |
| Hemoglobin (g/dL), mean (SD) | 8.7 (2.4) | 42 | 8.5 (1.7) | 25 | 0.85 |
| Packed cell volume (%), mean (SD) | 26 (6.8) | 42 | 26.3 (5.1) | 25 | 0.44 |
| No anaemia (hct > 33%), n (%) | 2 (4.8) | 42 | 2 (0.08) | 25 | |
| Mild anaemia (hct 25 - \| 33%), n (%) | 19 (45.2) | 42 | 12 (0.5) | 25 | |
| Moderate anaemia (hct 15 - \| 25%), n (%) | 21 (50) | 42 | 11 (0.4) | 25 | |
| Severe anaemia (hct ≤ 15%), n (%) | 0 (0.000) | 42 | 0 (0.000) | 25 | |
| **Micro-biology and other laboratory results on admission** | | | | | |

| | | | | | |
|---|---|---|---|---|---|
| HIV status positive, n (%) | 16 (38.1) | 42 | 9 (33.3) | 27 | 0.59 |
| Positive blood culture, n (%) | 29 (69) | 42 | 0 (0) | 27 | <0.0001 |
| WBC count ($10^3$/uL), mean (SD) | 23.9 (15.1) | 42 | 17 (7) | 25 | 0.68 |
| Neutrophil granulocytes ($10^3$/uL), mean (SD) | 16.2 (10.7) | 35 | 9.8 (6.6) | 22 | 0.02 |
| Plasmodium density (parasites/uL), geometric mean (SD) | 0 (0) | 42 | 0 (0) | 27 | |
| Malaria positive | 0 (0) | 42 | 0 (0) | 27 | |
| **Chest X-Ray results** | | | | | >0.99 |
| Normal X-ray, n (%) | 5 (13.5) | 37 | 4 (17.4) | 23 | |
| Other infiltrate/abnormality, n (%) | 2 (5.4) | 37 | 4 (17.4) | 23 | |
| Primary endpoint pneumonia, n (%) | 30 (81.1) | 37 | 15 (65.2) | 23 | |
| **Evolution during admission** | | | | | |
| Duration of admission (days): Median (IQR) | 5.1 (2.5-8.2) | 42 | 3.7 (2-5.1) | 27 | 0.18 |
| Case fatality rate (in hospital death) | 3 (7.1) | 42 | 0 (0) | 27 | 0.24 |

HCT= hematocrit; IQR= Interquartile Range; MUAC=middle upper arm circumference; SD= Standard Deviation; WAZ = weight-for-age Z score. † P-values for continuous variables were estimated through analysis of Mann-Whitney U test. P-values for categorical variables used Fisher's exact test.

**Table S2 Single etiology sample designations and distributions for training (n=120) and validation sets (n=47).**

167 blood samples (including 8 replicates) were assayed using the SOMAScan platform. Replicate samples were included in the training set but not in the validation set.

| Sample designation | counts | Training sample counts | Validation sample counts |
|---|---|---|---|
| Bacteria | 73* | 54 (45%) | 19 (40%) |
| Malaria | 43** | 28 (23%) | 15 (32%) |
| Virus | 51*** | 38 (32%) | 13 (28%) |
| **Total** | **167** | **120** (100%) | **47** (100%) |

* Bacteria samples include 31 Bacteria (culture-positive), 27 PCR-Bacteria (PCR-positive), and 11 Probable Bacteria samples (strong clinical evidence of bacterial infection including leukocytosis and radiographic endpoint pneumonia without direct culture or PCR evidence). Four samples, including 3 PCR-Bacteria and 1 Bacteria etiology, were repeated in the second SOMAScan assay.

** Malaria samples include 42 analyzed in the first SOMAScan assay and 1 repeated in the second SOMAScan assay.

*** Virus samples include 48 analyzed in the first SOMAScan assay and 3 repeated in the second SOMAScan assay.

**Table S3 Mixed infection sample designation sample distribution.**

In addition to the groups defined as single etiology per previous methods [2], several mixed infection groups were defined. These included the following classifications: bacteria and malaria, bacteria and virus, malaria and virus, and malaria and other. Mixed samples comprised a separate test set and were not used for model training or primary validation.

| **Sample designation** | **counts** |
|---|---|
| Control[1] | 13 |
| Repeat Valid control[2] | 3 |
| Bacteria & Malaria[3] | 9 |
| Virus & probable Bacterial secondary co-infection[4] | 11 |
| Malaria & Other[5] | 1 |
| Malaria & Virus[6] | 2 |
| **Total number (%)** | **39** (100%) |

1 Control samples were analyzed in the first SOMAScan assay

2 Repeat valid controls are control samples from the first SOMAScan assay that were repeated in the second SOMAScan assay

3 Bacteria & Malaria patients had confirmed malaria parasitemia according to the pre-specified age thresholds, but with a lobar consolidation on chest x-ray (“endpoint pneumonia”) or with a positive (not contaminated) blood culture; hence not fulfilling the case definition for either malaria or bacteria single etiology group.

4 Virus & probable secondary bacterial co-infection. Patients were confirmed viral positive (in the nasopharyngeal aspirate), were culture and PCR-negative for bacteria but with leukocytosis and radiographic endpoint pneumonia, suggestive of a secondary bacterial infection.

5 Malaria & Other patients had confirmed malaria parasitemia but with parasite density not fulfilling the pre-specified parasitemia age-thresholds (i.e. less than 2,500 asexual parasites/ml of blood in children over 1 year of age), or were patients with confirmed malaria parasitemia according to the pre-specified age-thresholds, but with infiltrates in the chest X ray and no evidence of positive blood culture, hence not not fulfilling the case definition for single etiology malaria.

6 Malaria & Virus patients fulfilled the same criteria as Malaria & Other, but were positive for virus in the nasopharyngeal aspirate.

**Table S4 List of markers for Bacteria vs Virus (n=219) and Bacteria vs Virus or Malaria (n=151).**

List of markers, selected using SAM [5] with FDR<0.01, and ranked using marker aggregation, for the two comparisons: Bacteria vs Virus and Bacteria vs Malaria or Virus, with 219 and 151 biomarkers, respectively. Each marker is assigned a score "score(d)" based on the difference (d) in gene expression, relative to the standard deviation of repeated measurements for that gene. Genes with scores greater than the threshold are deemed potentially significant. The q-value represents the FDR (in percent) and is an approximation (hence, when SAM is invoked with an FDR threshold of 0.01, some markers have a resulting q-value slightly larger than 1.00%; see documentation for the SAM function in [33]. The "fold-change" is the change of gene expression from one group (bacteria, virus, virus and malaria) to another.[5] The "score" for each marker is derived from the rank aggregation algorithm to determine a global consensus ranking for each marker ("final rank").

**(A)** All 219 protein markers for the Bacteria vs Virus comparison (FDR < 0.01)

| **Biomarker** | **Description** | **score(d)** | **fold change** | **q-value (%)** | **score** | **Final rank** |
|---|---|---|---|---|---|---|
| PLA2G2A | Phospholipase A2, membrane associated | -6.272 | 0.141 | 0 | 539 | 1 |
| LCN2 | Neutrophil gelatinase-associated lipocalin | -5.248 | 0.309 | 0 | 5629 | 2 |
| CCL23 | C-C motif chemokine 23 | -5.105 | 0.373 | 0 | 72528 | 3 |
| IL1RL1 | Interleukin-1 receptor-like 1 | -5.194 | 0.256 | 0 | 75304 | 4 |
| MPO | Myeloperoxidase | -4.75 | 0.56 | 0 | 122395 | 5 |
| CX3CL1 | Fractalkine | 4.145 | 1.679 | 0 | 178304 | 6 |
| RETN | Resistin | -4.542 | 0.477 | 0 | 210805 | 7 |

| | | | | | | |
|---|---|---|---|---|---|---|
| IL6 | Interleukin-6 | -4.062 | 0.248 | 0 | 372195 | 8 |
| NTN4 | Netrin-4 | -4.326 | 0.537 | 0 | 410795 | 9 |
| SERPINA4 | Kallistatin | 4.236 | 1.727 | 0 | 779067 | 10 |
| CSF3 | Granulocyte colony-stimulating factor | -3.019 | 0.446 | 0 | 809589 | 11 |
| CXCL13 | C-X-C motif chemokine 13 | -3.96 | 0.452 | 0 | 1144035 | 12 |
| CA6 | Carbonic anhydrase 6 | 4.052 | 4.145 | 0 | 1177340 | 13 |
| EPO | Erythropoietin | -3.093 | 0.474 | 0 | 1492008 | 14 |
| S100A9 | Protein S100-A9 | -3.726 | 0.575 | 0 | 2159784 | 15 |
| SERPINA1 | Alpha-1-antitrypsin | -4.197 | 0.512 | 0 | 2284000 | 16 |
| ITIH4 | Inter-alpha-trypsin inhibitor heavy chain H4 | -3.879 | 0.609 | 0 | 4599539 | 17 |
| HAMP | Hepcidin | -3.31 | 0.372 | 0 | 6823232 | 18 |
| FABP5 | Fatty acid-binding protein, epidermal | -3.707 | 0.619 | 0 | 7156875 | 19 |
| SLPI | Antileukoproteinase | -3.56 | 0.688 | 0 | 8165000 | 20 |
| SERPINA5 | Plasma serine protease inhibitor | 2.585 | 2.23 | 0.439 | 9812352 | 21 |
| PDE11A | Dual 3',5'-cyclic-AMP and -GMP phosphodiesterase 11A | -3.028 | 0.757 | 0 | 10829973 | 22 |
| CDC2.CCNB1 | Cyclin-dependent kinase 1:G2/mitotic-specific cyclin-B1 complex | -1.917 | 0.809 | 1.782 | 11202365 | 23 |
| PIK3CG | Phosphatidylinositol 4,5-bisphosphate 3-kinase catalytic subunit gamma isoform | -2.287 | 0.81 | 0.439 | 12178053 | 24 |
| CCL3 | C-C motif chemokine 3 | -2.792 | 0.686 | 0 | 12523072 | 25 |
| CTF1 | Cardiotrophin-1 | -2.281 | 0.81 | 0.439 | 12617096 | 26 |
| IMPDH1 | Inosine-5'-monophosphate dehydrogenase 1 | -3.597 | 0.603 | 0 | 13490256 | 27 |
| ESD | S-formylglutathione hydrolase | 3.23 | 1.683 | 0 | 16598579 | 28 |
| ROR1 | Tyrosine-protein kinase transmembrane receptor ROR1 | 3.597 | 1.627 | 0 | 17099712 | 29 |
| HGF | Hepatocyte growth factor | -3.82 | 0.506 | 0 | 17694336 | 30 |
| KLK8 | Kallikrein-8 | 3.337 | 1.895 | 0 | 21939160 | 31 |
| PAPPA | Pappalysin-1 | 3.129 | 1.89 | 0 | 22683136 | 32 |

| | | | | | | |
|---|---|---|---|---|---|---|
| SLITRK5 | SLIT and NTRK-like protein 5 | 3.166 | 2.011 | 0 | 23984160 | 33 |
| HIST1H1C | Histone H1.2 | -3.656 | 0.499 | 0 | 25727355 | 34 |
| CDON | Cell adhesion molecule-related/down-regulated by oncogenes | 3.442 | 2.177 | 0 | 27024015 | 35 |
| SHBG | Sex hormone-binding globulin | 3.267 | 1.735 | 0 | 42887296 | 36 |
| LTA.LTB.1 | Lymphotoxin alpha2:beta1 | 2.698 | 2.153 | 0 | 43073024 | 37 |
| TNFRSF21 | Tumor necrosis factor receptor superfamily member 21 | 3.657 | 1.56 | 0 | 46155057 | 38 |
| IL1R1 | Interleukin-1 receptor type 1 | 2.906 | 1.971 | 0 | 55924937 | 39 |
| HSD17B1 | Estradiol 17-beta-dehydrogenase 1 | -3.011 | 0.702 | 0 | 60892608 | 40 |
| MAPK1 | Mitogen-activated protein kinase 1 | -2.163 | 0.748 | 0.95 | 68465256 | 41 |
| CCL18 | C-C motif chemokine 18 | -3.314 | 0.569 | 0 | 70039553 | 42 |
| PGLYRP1 | Peptidoglycan recognition protein 1 | -2.713 | 0.451 | 0 | 83894265 | 43 |
| LTA.LTB | Lymphotoxin alpha1:beta2 | 3.218 | 1.866 | 0 | 85042953 | 44 |
| KIT | Mast/stem cell growth factor receptor Kit | 2.956 | 1.53 | 0 | 86318715 | 45 |
| AFM | Afamin | 3.643 | 1.512 | 0 | 87402621 | 46 |
| MATN3 | Matrilin-3 | -2.246 | 0.772 | 0.95 | 91152909 | 47 |
| Human.virus.2 | Protein Rev_HV2BE | -2.281 | 0.55 | 0.439 | 97576185 | 48 |
| MRC2 | C-type mannose receptor 2 | 3.373 | 1.843 | 0 | 99867485 | 49 |
| JAK2 | Tyrosine-protein kinase JAK2 | -3.075 | 0.649 | 0 | 99941040 | 50 |
| MST1R | Macrophage-stimulating protein receptor | -2.066 | 0.833 | 0.95 | 1.11E+08 | 51 |
| HP | Haptoglobin | -2.398 | 0.359 | 0.439 | 1.15E+08 | 52 |
| SERPINC1 | Antithrombin-III | 2.367 | 1.198 | 0.95 | 1.15E+08 | 53 |
| SPINT1 | Kunitz-type protease inhibitor 1 | 3.253 | 1.755 | 0 | 1.3E+08 | 54 |
| HBA1.HBB | Hemoglobin | 2.181 | 2.521 | 1.782 | 1.36E+08 | 55 |
| RB1 | Retinoblastoma-associated protein | -2.092 | 0.741 | 0.95 | 1.43E+08 | 56 |
| REN | Renin | -2.772 | 0.622 | 0 | 1.44E+08 | 57 |

| | | | | | | |
|---|---|---|---|---|---|---|
| CXCL1 | Growth-regulated alpha protein | 2.335 | 1.631 | 0.95 | 1.45E+08 | 58 |
| ERBB2 | Receptor tyrosine-protein kinase erbB-2 | -2.13 | 0.674 | 0.95 | 1.49E+08 | 59 |
| LTF | Lactotransferrin | -2.86 | 0.626 | 0 | 1.5E+08 | 60 |
| SAA1 | Serum amyloid A-1 protein | -2.474 | 0.355 | 0.439 | 1.5E+08 | 61 |
| LEP | Leptin | 2.313 | 1.51 | 0.95 | 1.51E+08 | 62 |
| ITGA2B.ITGB3 | Integrin alpha-IIb: beta-3 complex | 2.587 | 1.681 | 0.439 | 1.55E+08 | 63 |
| L1CAM | Neural cell adhesion molecule L1 | 2.699 | 1.49 | 0 | 1.6E+08 | 64 |
| CCL14 | C-C motif chemokine 14 | -2.707 | 0.67 | 0 | 1.61E+08 | 65 |
| CCL19 | C-C motif chemokine 19 | -2.404 | 0.664 | 0.439 | 1.64E+08 | 66 |
| MRC1 | Macrophage mannose receptor 1 | -3.052 | 0.664 | 0 | 1.64E+08 | 67 |
| CCL20 | C-C motif chemokine 20 | -2.226 | 0.722 | 0.95 | 1.66E+08 | 68 |
| HMGB1 | High mobility group protein B1 | -2.658 | 0.679 | 0 | 1.81E+08 | 69 |
| PLG.1 | Plasmin | -2.109 | 0.483 | 0.95 | 2.01E+08 | 70 |
| IGHD..IGK..IGL. | Immunoglobulin D | -1.923 | 0.508 | 1.782 | 2.09E+08 | 71 |
| CDH5 | Cadherin-5 | 3.08 | 1.349 | 0 | 2.13E+08 | 72 |
| CFHR5 | Complement factor H-related protein 5 | -2.644 | 0.609 | 0 | 2.15E+08 | 73 |
| CDH3 | Cadherin-3 | 2.605 | 1.363 | 0.439 | 2.2E+08 | 74 |
| CDK2.CCNA2 | Cyclin-dependent kinase 2:Cyclin-A2 complex | -2.558 | 0.678 | 0 | 2.3E+08 | 75 |
| PRSS1 | Trypsin-1 | -2.398 | 0.682 | 0.439 | 2.33E+08 | 76 |
| CNTN1 | Contactin-1 | 3.022 | 1.405 | 0 | 2.34E+08 | 77 |
| MET | Hepatocyte growth factor receptor | 2.137 | 1.24 | 1.782 | 2.45E+08 | 78 |
| NTRK2 | BDNF/NT-3 growth factors receptor | 3.142 | 1.522 | 0 | 2.48E+08 | 79 |
| NCAM1 | Neural cell adhesion molecule 1, 120 kDa isoform | 2.732 | 1.438 | 0 | 2.54E+08 | 80 |
| PRSS2 | Trypsin-2 | -2.696 | 0.519 | 0 | 2.54E+08 | 81 |
| SERPINE1 | Plasminogen activator inhibitor 1 | -2.126 | 0.594 | 0.95 | 2.58E+08 | 82 |
| FGA.FGB. | D-dimer | -2.435 | 0.529 | 0.439 | 2.75E+08 | 83 |

| | | | | | | |
|---|---|---|---|---|---|---|
| FGG.1 | | | | | | |
| PPID | Peptidyl-prolyl cis-trans isomerase D | -2.662 | 0.628 | 0 | 2.78E+08 | 84 |
| XPNPEP1 | Xaa-Pro aminopeptidase 1 | 2.467 | 1.557 | 0.439 | 2.96E+08 | 85 |
| CADM1 | Cell adhesion molecule 1 | 2.802 | 1.579 | 0 | 2.99E+08 | 86 |
| LSAMP | Limbic system-associated membrane protein | 2.591 | 1.338 | 0.439 | 3.2E+08 | 87 |
| UNC5D | Netrin receptor UNC5D | 2.613 | 1.758 | 0.439 | 3.22E+08 | 88 |
| FGF23 | Fibroblast growth factor 23 | -1.985 | 0.666 | 1.782 | 3.23E+08 | 89 |
| ZAP70 | Tyrosine-protein kinase ZAP-70 | -2.013 | 0.825 | 1.782 | 3.24E+08 | 90 |
| NTRK3 | NT-3 growth factor receptor | 2.727 | 1.517 | 0 | 3.28E+08 | 91 |
| PLXNC1 | Plexin-C1 | 2.249 | 1.248 | 0.95 | 3.29E+08 | 92 |
| PRTN3 | Myeloblastin | -3.314 | 0.599 | 0 | 3.41E+08 | 93 |
| CCL13 | C-C motif chemokine 13 | -2.471 | 0.685 | 0.439 | 3.52E+08 | 94 |
| TFPI | Tissue factor pathway inhibitor | -3.09 | 0.61 | 0 | 3.54E+08 | 95 |
| TNFRSF25 | Tumor necrosis factor receptor superfamily member 25 | -2.227 | 0.738 | 0.95 | 3.57E+08 | 96 |
| ETHE1 | Persulfide dioxygenase ETHE1, mitochondrial | -2.449 | 0.762 | 0.439 | 3.62E+08 | 97 |
| TPSB2 | Tryptase beta-2 | 2.201 | 1.512 | 1.782 | 3.68E+08 | 98 |
| CCL7 | C-C motif chemokine 7 | 2.398 | 1.7 | 0.439 | 3.71E+08 | 99 |
| KLK7 | Kallikrein-7 | 2.36 | 1.632 | 0.95 | 3.72E+08 | 100 |
| H2AFZ | Histone H2A.z | -2.385 | 0.548 | 0.439 | 3.73E+08 | 101 |
| RPS7 | 40S ribosomal protein S7 | -2.058 | 0.701 | 0.95 | 3.96E+08 | 102 |
| IGFBP3 | Insulin-like growth factor-binding protein 3 | 3.114 | 1.507 | 0 | 3.96E+08 | 103 |
| RGMB | RGM domain family member B | 2.153 | 1.207 | 1.782 | 3.97E+08 | 104 |
| CHIT1 | Chitotriosidase-1 | -2.205 | 0.655 | 0.95 | 4.01E+08 | 105 |
| STK17B | Serine/threonine-protein kinase 17B | -2.195 | 0.772 | 0.95 | 4.03E+08 | 106 |
| CXCL3.CXCL2 | Gro-beta/gamma | 2.133 | 1.647 | 1.782 | 4.07E+08 | 107 |

| | | | | | | |
|---|---|---|---|---|---|---|
| PECAM1 | Platelet endothelial cell adhesion molecule | -2.097 | 0.819 | 0.95 | 4.08E+08 | 108 |
| CNTFR | Ciliary neurotrophic factor receptor subunit alpha | 2.445 | 1.427 | 0.439 | 4.14E+08 | 109 |
| EPHB2 | Ephrin type-B receptor 2 | 3.234 | 1.517 | 0 | 4.16E+08 | 110 |
| IL12A.IL12B | Interleukin-12 | -2.444 | 0.701 | 0.439 | 4.21E+08 | 111 |
| HSPA8 | Heat shock cognate 71 kDa protein | -2.042 | 0.639 | 0.95 | 4.25E+08 | 112 |
| HAPLN1 | Hyaluronan and proteoglycan link protein 1 | 2.503 | 1.65 | 0.439 | 4.27E+08 | 113 |
| SHH | Sonic hedgehog protein | 2.615 | 1.56 | 0.439 | 4.32E+08 | 115 |
| ICOSLG | ICOS ligand | -2.206 | 0.702 | 0.95 | 4.37E+08 | 116 |
| OCIAD1 | OCIA domain-containing protein 1 | -2.162 | 0.688 | 0.95 | 4.38E+08 | 117 |
| EPHA2 | Ephrin type-A receptor 2 | 2.524 | 1.418 | 0.439 | 4.42E+08 | 118 |
| RBM39 | RNA-binding protein 39 | -2.391 | 0.752 | 0.439 | 4.42E+08 | 119 |
| EPHA5 | Ephrin type-A receptor 5 | 2.745 | 1.508 | 0 | 4.5E+08 | 120 |
| TNFRSF1A | Tumor necrosis factor receptor superfamily member 1A | -2.746 | 0.708 | 0 | 4.51E+08 | 121 |
| SFTPD | Pulmonary surfactant-associated protein D | -2.507 | 0.642 | 0 | 4.73E+08 | 122 |
| CD109 | CD109 antigen | 2.715 | 1.589 | 0 | 4.74E+08 | 123 |
| ANGPT2 | Angiopoietin-2 | -2.212 | 0.766 | 0.95 | 4.77E+08 | 124 |
| LRP8 | Low-density lipoprotein receptor-related protein 8 | 2.561 | 1.367 | 0.439 | 4.81E+08 | 125 |
| DKK3 | Dickkopf-related protein 3 | 2.647 | 1.407 | 0 | 4.82E+08 | 126 |
| ACAN | Aggrecan core protein | 2.322 | 1.508 | 0.95 | 4.83E+08 | 127 |
| KLK14 | Kallikrein-14 | -2.281 | 0.818 | 0.439 | 4.89E+08 | 128 |
| FGFR1 | Fibroblast growth factor receptor 1 | 3.237 | 1.451 | 0 | 4.89E+08 | 129 |
| PRKCZ | Protein kinase C zeta type | -2.524 | 0.731 | 0 | 4.98E+08 | 130 |
| LRIG3 | Leucine-rich repeats and immunoglobulin-like domains protein 3 | 2.409 | 1.373 | 0.439 | 4.98E+08 | 131 |
| ADCYAP1 | Pituitary adenylate cyclase-activating | -2.532 | 0.652 | 0 | 5E+08 | 132 |

| | | | | | | |
|---|---|---|---|---|---|---|
| .1 | polypeptide 38 | | | | | |
| CAST | Calpastatin | 2.848 | 1.526 | 0 | 5.01E+08 | 133 |
| PGD | 6-phosphogluconate dehydrogenase, decarboxylating | -1.967 | 0.652 | 1.782 | 5.02E+08 | 134 |
| TOP1 | DNA topoisomerase 1 | -2 | 0.704 | 1.782 | 5.05E+08 | 135 |
| MAPKAP K3 | MAP kinase-activated protein kinase 3 | -2.578 | 0.731 | 0 | 5.08E+08 | 136 |
| CHL1 | Neural cell adhesion molecule L1-like protein | 2.582 | 1.41 | 0.439 | 5.1E+08 | 137 |
| DSG2 | Desmoglein-2 | -2.013 | 0.762 | 1.782 | 5.12E+08 | 138 |
| BIRC7 | Baculoviral IAP repeat-containing protein 7 Isoform beta | -2.105 | 0.805 | 0.95 | 5.21E+08 | 139 |
| VEGFA | Vascular endothelial growth factor A | -2.076 | 0.813 | 0.95 | 5.24E+08 | 140 |
| EPHB6 | Ephrin type-B receptor 6 | 2.591 | 1.32 | 0.439 | 5.26E+08 | 141 |
| NOTCH1 | Neurogenic locus notch homolog protein 1 | 2.251 | 1.195 | 0.95 | 5.28E+08 | 142 |
| LPO | Lactoperoxidase | 2.303 | 1.765 | 0.95 | 5.38E+08 | 143 |
| GSN | Gelsolin | 2.851 | 1.33 | 0 | 5.4E+08 | 144 |
| LILRB2 | Leukocyte immunoglobulin-like receptor subfamily B member 2 | -1.895 | 0.755 | 1.782 | 5.6E+08 | 145 |
| FLT3 | Receptor-type tyrosine-protein kinase FLT3 | -2.032 | 0.754 | 0.95 | 5.76E+08 | 146 |
| EFNA5 | Ephrin-A5 | 2.569 | 1.384 | 0.439 | 5.84E+08 | 147 |
| BPI | Bactericidal permeability-increasing protein | -2.817 | 0.57 | 0 | 5.91E+08 | 148 |
| FCN2 | Ficolin-2 | -2.557 | 0.724 | 0 | 5.96E+08 | 149 |
| SIRT2 | NAD-dependent protein deacetylase sirtuin-2 | -2.177 | 0.727 | 0.95 | 6E+08 | 150 |
| FSTL3 | Follistatin-related protein 3 | -1.978 | 0.768 | 1.782 | 6.01E+08 | 151 |
| PDCD1LG 2 | Programmed cell death 1 ligand 2 | 2.373 | 1.318 | 0.95 | 6.03E+08 | 152 |
| CXCL12 | Stromal cell-derived factor 1 | -2.679 | 0.639 | 0 | 6.09E+08 | 153 |

| | | | | | | |
|---|---|---|---|---|---|---|
| CNTN2 | Contactin-2 | 2.308 | 1.37 | 0.95 | 6.27E+08 | 154 |
| TDGF1 | Teratocarcinoma-derived growth factor 1 | -2.168 | 0.764 | 0.95 | 6.39E+08 | 155 |
| CD200 | OX-2 membrane glycoprotein | 2.364 | 1.308 | 0.95 | 6.48E+08 | 156 |
| ALPL | Alkaline phosphatase, tissue-nonspecific isozyme | -2.578 | 0.711 | 0 | 6.91E+08 | 157 |
| FGF12 | Fibroblast growth factor 12 | -2.017 | 0.804 | 1.782 | 6.91E+08 | 158 |
| FN1.2 | Fibronectin | 2.136 | 1.625 | 1.782 | 6.93E+08 | 159 |
| CST7 | Cystatin-F | -2.045 | 0.646 | 0.95 | 6.96E+08 | 160 |
| IL22RA1 | Interleukin-22 receptor subunit alpha-1 | -2.092 | 0.729 | 0.95 | 7.08E+08 | 161 |
| SEMA3A | Semaphorin-3A | -2.378 | 0.716 | 0.439 | 7.19E+08 | 162 |
| IGFBP2 | Insulin-like growth factor-binding protein 2 | -2.777 | 0.674 | 0 | 7.3E+08 | 163 |
| GFRA1 | GDNF family receptor alpha-1 | -2.236 | 0.728 | 0.95 | 7.46E+08 | 164 |
| F9.1 | Coagulation factor IXab | -2.335 | 0.786 | 0.439 | 7.77E+08 | 165 |
| KAT6A | Histone acetyltransferase KAT6A | -2.228 | 0.793 | 0.95 | 7.94E+08 | 166 |
| HSP90AA1.HSP90AB1 | Heat shock protein HSP 90-alpha/beta | -1.933 | 0.774 | 1.782 | 8.31E+08 | 167 |
| ADRBK1 | beta-adrenergic receptor kinase 1 | -2.635 | 0.73 | 0 | 8.36E+08 | 168 |
| RTN4 | Reticulon-4 | -2.057 | 0.767 | 0.95 | 8.77E+08 | 169 |
| DAPK2 | Death-associated protein kinase 2 | -2.298 | 0.78 | 0.439 | 8.86E+08 | 170 |
| MMEL1 | Membrane metallo-endopeptidase-like 1 | -2.221 | 0.78 | 0.95 | 8.87E+08 | 171 |
| OMD | Osteomodulin | 2.11 | 1.368 | 1.782 | 8.92E+08 | 172 |
| IL18R1 | Interleukin-18 receptor 1 | -2.022 | 0.779 | 0.95 | 8.96E+08 | 173 |
| EPHA1 | Ephrin type-A receptor 1 | 2.198 | 1.577 | 1.782 | 9.36E+08 | 174 |
| SERPING1 | Plasma protease C1 inhibitor | 2.582 | 1.413 | 0.439 | 9.38E+08 | 175 |
| ALCAM | CD166 antigen | 2.52 | 1.27 | 0.439 | 9.38E+08 | 176 |
| DKKL1 | Dickkopf-like protein 1 | -2.283 | 0.795 | 0.439 | 9.81E+08 | 177 |
| AHSG | Alpha-2-HS-glycoprotein | 2.568 | 1.351 | 0.439 | 9.96E+08 | 178 |
| CCL3L1 | C-C motif chemokine 3-like 1 | -2.289 | 0.778 | 0.439 | 1.02E+09 | 179 |

| | | | | | | |
|---|---|---|---|---|---|---|
| NOTCH3 | Neurogenic locus notch homolog protein 3 | 2.141 | 1.275 | 1.782 | 1.03E+09 | 180 |
| HDGFRP2 | Hepatoma-derived growth factor-related protein 2 | -2.38 | 0.769 | 0.439 | 1.05E+09 | 181 |
| MMP17 | Matrix metalloproteinase-17 | -2.42 | 0.732 | 0.439 | 1.05E+09 | 182 |
| PPP3R1 | Calcineurin subunit B type 1 | -2.257 | 0.804 | 0.95 | 1.07E+09 | 183 |
| SLAMF7 | SLAM family member 7 | -1.928 | 0.776 | 1.782 | 1.07E+09 | 184 |
| UNC5C | Netrin receptor UNC5C | 2.245 | 1.249 | 0.95 | 1.08E+09 | 185 |
| ANGPTL4 | Angiopoietin-related protein 4 | -2.32 | 0.714 | 0.439 | 1.1E+09 | 186 |
| DPT | Dermatopontin | 2.21 | 1.449 | 0.95 | 1.1E+09 | 187 |
| F10 | Coagulation factor Xa | -2.252 | 0.805 | 0.95 | 1.11E+09 | 188 |
| PDE2A | cGMP-dependent 3',5'-cyclic phosphodiesterase | -2.017 | 0.802 | 1.782 | 1.11E+09 | 189 |
| CTSG | Cathepsin G | -1.993 | 0.766 | 1.782 | 1.12E+09 | 190 |
| WISP1 | WNT1-inducible-signaling pathway protein 1 | -2.119 | 0.8 | 0.95 | 1.12E+09 | 191 |
| LCMT1 | Leucine carboxyl methyltransferase 1 | -2.108 | 0.785 | 0.95 | 1.12E+09 | 192 |
| KLKB1 | Plasma kallikrein | 2.182 | 1.267 | 1.782 | 1.13E+09 | 193 |
| LBP | Lipopolysaccharide-binding protein | -1.975 | 0.71 | 1.782 | 1.13E+09 | 194 |
| IL17D | Interleukin-17D | -1.964 | 0.819 | 1.782 | 1.15E+09 | 195 |
| SMPDL3A | Acid sphingomyelinase-like phosphodiesterase 3a | -2.038 | 0.778 | 0.95 | 1.15E+09 | 196 |
| BCAN | Brevican core protein | 2.418 | 1.494 | 0.439 | 1.15E+09 | 197 |
| PRKCD | Protein kinase C delta type | -2.138 | 0.806 | 0.95 | 1.17E+09 | 198 |
| CNTN4 | Contactin-4 | 2.227 | 1.286 | 0.95 | 1.18E+09 | 199 |
| THBS4 | Thrombospondin-4 | 2.446 | 1.45 | 0.439 | 1.2E+09 | 200 |
| NRXN1 | Neurexin-1-beta | 2.38 | 1.446 | 0.95 | 1.24E+09 | 201 |
| ANP32B | Acidic leucine-rich nuclear phosphoprotein 32 family member B | -1.974 | 0.822 | 1.782 | 1.25E+09 | 202 |
| ADCYAP1 | Pituitary adenylate cyclase-activating polypeptide 27 | -2.2 | 0.72 | 0.95 | 1.25E+09 | 203 |

| | | | | | | |
|---|---|---|---|---|---|---|
| IFNL2 | Interferon lambda-2 | -1.899 | 0.822 | 1.782 | 1.26E+09 | 204 |
| WIF1 | Wnt inhibitory factor 1 | 2.235 | 1.363 | 0.95 | 1.33E+09 | 205 |
| F10.1 | Coagulation Factor X | -1.891 | 0.838 | 1.782 | 1.34E+09 | 206 |
| TIMP1 | Metalloproteinase inhibitor 1 | -2.037 | 0.709 | 0.95 | 1.35E+09 | 207 |
| CKM | Creatine kinase M-type | 2.274 | 1.467 | 0.95 | 1.37E+09 | 208 |
| MDM2 | E3 ubiquitin-protein ligase Mdm2 | -2.336 | 0.815 | 0.439 | 1.48E+09 | 209 |
| AGT | Angiotensinogen | -1.895 | 0.762 | 1.782 | 1.48E+09 | 210 |
| CASP10 | Caspase-10 | -1.901 | 0.866 | 1.782 | 1.49E+09 | 211 |
| MAPK14 | Mitogen-activated protein kinase 14 | -2.062 | 0.801 | 0.95 | 1.5E+09 | 212 |
| CRISP3 | Cysteine-rich secretory protein 3 | -1.914 | 0.836 | 1.782 | 1.51E+09 | 213 |
| STC1 | Stanniocalcin-1 | -1.972 | 0.768 | 1.782 | 1.57E+09 | 214 |
| GFRA3 | GDNF family receptor alpha-3 | -1.911 | 0.779 | 1.782 | 1.64E+09 | 215 |
| RSPO2 | R-spondin-2 | -1.91 | 0.787 | 1.782 | 1.86E+09 | 216 |
| F9 | Coagulation factor IX | -1.958 | 0.835 | 1.782 | 1.87E+09 | 217 |
| APOD | Apolipoprotein D | -2.05 | 0.833 | 0.95 | 1.88E+09 | 218 |
| CCL27 | C-C motif chemokine 27 | -2.15 | 0.8 | 0.95 | 1.93E+09 | 219 |

**(B)** All 151 protein markers for the Bacteria vs Virus or Malaria comparison (FDR < 0.01)

| Biomarker | Description | score(d) | fold change | q-value (%) | score | Final rank |
|---|---|---|---|---|---|---|
| HP | Haptoglobin | -4.624 | 0.097 | 0 | 3817 | 1 |
| CX3CL1 | Fractalkine | 4.894 | 1.752 | 0 | 4217 | 2 |
| RETN | Resistin | -4.98 | 0.512 | 0 | 7053 | 3 |
| CSF3 | Granulocyte colony-stimulating factor | -4.377 | 0.386 | 0 | 27321 | 4 |
| LCN2 | Neutrophil gelatinase-associated lipocalin | -5.398 | 0.367 | 0 | 30544 | 5 |
| SLPI | Antileukoproteinase | -4.457 | 0.66 | 0 | 79632 | 6 |
| ESD | S-formylglutathione hydrolase | 4.17 | 1.78 | 0 | 103631 | 7 |
| KYNU | Kynureninase | 3.819 | 1.786 | 0 | 207560 | 8 |
| IL6 | Interleukin-6 | -4.184 | 0.304 | 0 | 240832 | 9 |
| S100A9 | Protein S100-A9 | -4.857 | 0.543 | 0 | 250112 | 10 |
| HBA1.HBB | Hemoglobin | 4.248 | 7.32 | 0 | 332432 | 11 |
| PLA2G2A | Phospholipase A2, membrane associated | -3.912 | 0.309 | 0 | 443185 | 12 |
| SERPINA4 | Kallistatin | 4.253 | 1.571 | 0 | 507115 | 13 |
| PIK3CG | Phosphatidylinositol 4,5-bisphosphate 3-kinase catalytic subunit gamma isoform | -3.155 | 0.783 | 0 | 631496 | 14 |
| CXCL13 | C-X-C motif chemokine 13 | -3.886 | 0.516 | 0 | 671021 | 15 |
| FABP5 | Fatty acid-binding protein, epidermal | -4.271 | 0.626 | 0 | 770688 | 16 |
| CD36 | Platelet glycoprotein 4 | 3.768 | 2.829 | 0 | 906752 | 17 |
| PAPPA | Pappalysin-1 | 3.399 | 1.825 | 0 | 2123200 | 18 |
| FGF17 | Fibroblast growth factor 17 | -2.456 | 0.775 | 0.619 | 2471135 | 19 |
| HGF | Hepatocyte growth factor | -4.516 | 0.523 | 0 | 2516560 | 20 |
| GZMB | Granzyme B | 3.107 | 1.933 | 0 | 3226520 | 21 |
| NAGK | N-acetyl-D-glucosamine kinase | 3.556 | 1.641 | 0 | 3750264 | 22 |
| CD274 | Programmed cell death 1 ligand 1 | 2.777 | 1.422 | 0 | 5022381 | 23 |

| | | | | | | |
|---|---|---|---|---|---|---|
| IL1RL1 | Interleukin-1 receptor-like 1 | -3.371 | 0.433 | 0 | 5061257 | 24 |
| LTA.LTB.1 | Lymphotoxin alpha2:beta1 | 3.274 | 2.046 | 0 | 7261947 | 25 |
| BIRC7 | Baculoviral IAP repeat-containing protein 7 Isoform beta | -3.29 | 0.756 | 0 | 7726488 | 26 |
| MPO | Myeloperoxidase | -3.561 | 0.669 | 0 | 8970291 | 27 |
| LAG3 | Lymphocyte activation gene 3 protein | 3.672 | 1.958 | 0 | 9707152 | 28 |
| FGA.FGB.FGG.1 | D-dimer | -3.351 | 0.501 | 0 | 12127520 | 29 |
| CTF1 | Cardiotrophin-1 | -2.757 | 0.796 | 0 | 12407197 | 30 |
| HAMP | Hepcidin | -3.041 | 0.429 | 0 | 12572847 | 31 |
| CFHR5 | Complement factor H-related protein 5 | -3.044 | 0.62 | 0 | 14080573 | 32 |
| ETHE1 | Persulfide dioxygenase ETHE1, mitochondrial | -3.282 | 0.73 | 0 | 14145024 | 33 |
| CCL21 | C-C motif chemokine 21 | -3.064 | 0.7 | 0 | 15049216 | 34 |
| IGFBP3 | Insulin-like growth factor-binding protein 3 | 3.235 | 1.449 | 0 | 15969024 | 35 |
| HSD17B1 | Estradiol 17-beta-dehydrogenase 1 | -2.573 | 0.764 | 0 | 17290496 | 36 |
| PLG.1 | Plasmin | -2.847 | 0.471 | 0 | 17838352 | 37 |
| PTPN11 | Tyrosine-protein phosphatase non-receptor type 11 | 3.081 | 1.445 | 0 | 18420931 | 38 |
| RB1 | Retinoblastoma-associated protein | -2.135 | 0.774 | 0.994 | 18433605 | 39 |
| MST1R | Macrophage-stimulating protein receptor | -2.643 | 0.823 | 0 | 20582656 | 40 |
| PRDX6 | Peroxiredoxin-6 | 2.924 | 1.651 | 0 | 21192460 | 41 |
| Human.virus.2 | Protein Rev_HV2BE | -3.175 | 0.527 | 0 | 21518796 | 42 |
| C4A.C4B.1 | Complement C4 | 2.634 | 3.126 | 0.619 | 22014388 | 43 |
| CCL3 | C-C motif chemokine 3 | -2.604 | 0.747 | 0 | 22561200 | 44 |
| F2.1 | Prothrombin | 2.744 | 2.463 | 0 | 23337369 | 45 |
| CCL7 | C-C motif chemokine 7 | 2.641 | 1.625 | 0.619 | 26251264 | 46 |
| HSPA8 | Heat shock cognate 71 kDa protein | -3.295 | 0.565 | 0 | 29458053 | 47 |

| | | | | | | |
|---|---|---|---|---|---|---|
| XPNPEP1 | Xaa-Pro aminopeptidase 1 | 2.926 | 1.581 | 0 | 35900928 | 48 |
| JAK2 | Tyrosine-protein kinase JAK2 | -2.951 | 0.696 | 0 | 37850112 | 49 |
| ITIH4 | Inter-alpha-trypsin inhibitor heavy chain H4 | -3.128 | 0.699 | 0 | 39478731 | 50 |
| PGLYRP1 | Peptidoglycan recognition protein 1 | -2.687 | 0.499 | 0 | 43464008 | 51 |
| PRSS2 | Trypsin-2 | -2.621 | 0.579 | 0 | 47073468 | 52 |
| FN1.1 | Fibronectin Fragment 4 | 2.51 | 2.088 | 0.994 | 49025934 | 53 |
| ERBB2 | Receptor tyrosine-protein kinase erbB-2 | -2.655 | 0.672 | 0 | 50269135 | 54 |
| PLXNC1 | Plexin-C1 | 2.917 | 1.298 | 0 | 51597600 | 55 |
| CCL23 | C-C motif chemokine 23 | -3.052 | 0.572 | 0 | 51770916 | 56 |
| KIT | Mast/stem cell growth factor receptor Kit | 2.433 | 1.346 | 1.541 | 55683456 | 57 |
| CST7 | Cystatin-F | -3.28 | 0.568 | 0 | 56863744 | 58 |
| ADCYAP1.1 | Pituitary adenylate cyclase-activating polypeptide 38 | -2.695 | 0.692 | 0 | 57811648 | 59 |
| SAA1 | Serum amyloid A-1 protein | -2.3 | 0.443 | 0.994 | 57814700 | 60 |
| GZMA | Granzyme A | 3.127 | 1.715 | 0 | 57857019 | 61 |
| IL12B.IL23A | Interleukin-23 | -2.654 | 0.702 | 0 | 62266512 | 62 |
| CXCL6 | C-X-C motif chemokine 6 | 2.445 | 1.528 | 0.994 | 62403279 | 63 |
| ANGPT2 | Angiopoietin-2 | -2.801 | 0.761 | 0 | 68371275 | 64 |
| AMH | Muellerian-inhibiting factor | -2.68 | 0.662 | 0 | 68765655 | 65 |
| PDE11A | Dual 3',5'-cyclic-AMP and -GMP phosphodiesterase 11A | -2.224 | 0.834 | 0.994 | 71374451 | 66 |
| MMP12 | Macrophage metalloelastase | 2.603 | 1.72 | 0.619 | 71629448 | 67 |
| CCL19 | C-C motif chemokine 19 | -2.543 | 0.701 | 0 | 75809104 | 68 |
| REN | Renin | -2.526 | 0.697 | 0 | 77300480 | 69 |
| CCL14 | C-C motif chemokine 14 | -3.021 | 0.684 | 0 | 77632147 | 70 |
| IL18RAP | Interleukin-18 receptor accessory protein | -2.256 | 0.864 | 0.994 | 80479952 | 71 |
| VCAM1 | Vascular cell adhesion protein 1 | 3.265 | 1.462 | 0 | 80621568 | 72 |
| LRP8 | Low-density lipoprotein receptor-related | 2.41 | 1.274 | 1.541 | 84357630 | 73 |

| | | | | | | |
|---|---|---|---|---|---|---|
| | protein 8 | | | | | |
| NTN4 | Netrin-4 | -2.856 | 0.689 | 0 | 84643328 | 74 |
| LDHB | L-lactate dehydrogenase B chain | 2.999 | 1.383 | 0 | 91042368 | 75 |
| BIRC5 | Baculoviral IAP repeat-containing protein 5 | -2.512 | 0.831 | 0 | 92105445 | 76 |
| MDH1 | Malate dehydrogenase, cytoplasmic | 2.613 | 1.355 | 0.619 | 93004548 | 77 |
| MATN3 | Matrilin-3 | -2.308 | 0.798 | 0.994 | 93669087 | 78 |
| CNTN1 | Contactin-1 | 2.87 | 1.32 | 0 | 94727345 | 79 |
| SEMA3A | Semaphorin-3A | -2.943 | 0.703 | 0 | 96847465 | 80 |
| CCL28 | C-C motif chemokine 28 | -2.128 | 0.754 | 1.541 | 97300272 | 81 |
| HPX | Hemopexin | -2.237 | 0.702 | 0.994 | 100470051 | 82 |
| CAT | Catalase | 2.615 | 1.472 | 0.619 | 100571087 | 83 |
| CHIT1 | Chitotriosidase-1 | -2.416 | 0.669 | 0.619 | 100753484 | 84 |
| CD48 | CD48 antigen | 2.811 | 1.336 | 0 | 100837845 | 85 |
| SPINT1 | Kunitz-type protease inhibitor 1 | 2.426 | 1.429 | 1.541 | 101040423 | 86 |
| GSN | Gelsolin | 2.456 | 1.236 | 0.994 | 105135723 | 87 |
| EHMT2 | Histone-lysine N-methyltransferase EHMT2 | 2.443 | 1.317 | 0.994 | 108038980 | 88 |
| FGF18 | Fibroblast growth factor 18 | -2.156 | 0.701 | 0.994 | 108245676 | 89 |
| CSF1R | Macrophage colony-stimulating factor 1 receptor | 3.206 | 1.508 | 0 | 109648316 | 90 |
| FN1.2 | Fibronectin | 2.5 | 1.59 | 0.994 | 110208276 | 91 |
| SHBG | Sex hormone-binding globulin | 3.526 | 1.662 | 0 | 112985616 | 92 |
| HIST1H1C | Histone H1.2 | -2.454 | 0.657 | 0.619 | 118866217 | 93 |
| C2 | Complement C2 | -2.289 | 0.745 | 0.994 | 120470336 | 94 |
| SERPINA1 | Alpha-1-antitrypsin | -3.066 | 0.651 | 0 | 122448161 | 95 |
| C3.2 | C3a anaphylatoxin des Arginine | 2.425 | 1.682 | 1.541 | 123565862 | 96 |
| IL18R1 | Interleukin-18 receptor 1 | -2.843 | 0.747 | 0 | 124216768 | 97 |
| VEGFA | Vascular endothelial growth factor A | -2.701 | 0.781 | 0 | 130621440 | 98 |

| | | | | | | |
|---|---|---|---|---|---|---|
| CDH5 | Cadherin-5 | 2.825 | 1.265 | 0 | 132077225 | 99 |
| B2M | Beta-2-microglobulin | 2.398 | 1.282 | 1.541 | 141846650 | 100 |
| OCIAD1 | OCIA domain-containing protein 1 | -2.164 | 0.742 | 0.994 | 142408170 | 101 |
| PDCD1LG 2 | Programmed cell death 1 ligand 2 | 2.701 | 1.298 | 0 | 147354424 | 102 |
| IGFBP2 | Insulin-like growth factor-binding protein 2 | -2.373 | 0.743 | 0.619 | 155397250 | 103 |
| MMP7 | Matrilysin | -2.153 | 0.759 | 0.994 | 155548796 | 104 |
| VEGFA.1 | Vascular endothelial growth factor A, isoform 121 | -2.375 | 0.57 | 0.619 | 167420205 | 105 |
| CRLF1.CL CF1 | Cytokine receptor-like factor 1:Cardiotrophin-like cytokine factor 1 Complex | -2.582 | 0.786 | 0 | 177490310 | 106 |
| ABL1 | Tyrosine-protein kinase ABL1 | -2.382 | 0.759 | 0.619 | 178015814 | 107 |
| PROS1 | Vitamin K-dependent protein S | -2.282 | 0.84 | 0.994 | 180593480 | 108 |
| AFM | Afamin | 2.915 | 1.348 | 0 | 188496175 | 109 |
| ECE1 | Endothelin-converting enzyme 1 | -2.096 | 0.767 | 1.541 | 193310813 | 110 |
| MMP8 | Neutrophil collagenase | -2.136 | 0.657 | 0.994 | 194324063 | 111 |
| PON1 | Serum paraoxonase/arylesterase 1 | -2.119 | 0.778 | 1.541 | 199967425 | 112 |
| PTPN6 | Tyrosine-protein phosphatase non-receptor type 6 | 2.402 | 1.302 | 1.541 | 200334821 | 113 |
| TFPI | Tissue factor pathway inhibitor | -2.684 | 0.69 | 0 | 203267988 | 114 |
| FGFR1 | Fibroblast growth factor receptor 1 | 2.384 | 1.272 | 1.541 | 205188228 | 115 |
| PRSS1 | Trypsin-1 | -2.464 | 0.706 | 0.619 | 214221360 | 116 |
| TGFB3 | Transforming growth factor beta-3 | -2.151 | 0.785 | 0.994 | 215796832 | 117 |
| ALPL | Alkaline phosphatase, tissue-nonspecific isozyme | -2.283 | 0.759 | 0.994 | 225378604 | 118 |
| AHSG | Alpha-2-HS-glycoprotein | 2.52 | 1.288 | 0.994 | 237444991 | 119 |
| IL12A.IL1 2B | Interleukin-12 | -2.12 | 0.774 | 1.541 | 238871625 | 120 |
| LTF | Lactotransferrin | -2.401 | 0.698 | 0.619 | 244290928 | 121 |
| NXPH1 | Neurexophilin-1 | -2.108 | 0.786 | 1.541 | 256912273 | 122 |

| | | | | | | |
|---|---|---|---|---|---|---|
| MRC2 | C-type mannose receptor 2 | 2.548 | 1.487 | 0.994 | 258936128 | 123 |
| EPHA5 | Ephrin type-A receptor 5 | 2.716 | 1.426 | 0 | 261296460 | 124 |
| IL17A | Interleukin-17A | -2.207 | 0.795 | 0.994 | 265322596 | 125 |
| FCN2 | Ficolin-2 | -2.219 | 0.789 | 0.994 | 273131276 | 126 |
| GFRA1 | GDNF family receptor alpha-1 | -2.437 | 0.744 | 0.619 | 280357585 | 127 |
| CXCL12 | Stromal cell-derived factor 1 | -2.353 | 0.72 | 0.619 | 283038112 | 128 |
| HINT1 | Histidine triad nucleotide-binding protein 1 | -2.128 | 0.787 | 1.541 | 284657196 | 129 |
| TXNDC12 | Thioredoxin domain-containing protein 12 | -2.157 | 0.777 | 0.994 | 289318896 | 130 |
| CCL3L1 | C-C motif chemokine 3-like 1 | -2.233 | 0.809 | 0.994 | 295039826 | 131 |
| PRKCD | Protein kinase C delta type | -2.461 | 0.817 | 0.619 | 317039067 | 132 |
| SPON1 | Spondin-1 | -2.108 | 0.8 | 1.541 | 320281250 | 133 |
| ZAP70 | Tyrosine-protein kinase ZAP-70 | -2.271 | 0.825 | 0.994 | 322273663 | 134 |
| APCS | Serum amyloid P-component | -2.563 | 0.806 | 0 | 331490535 | 135 |
| CD97 | CD97 antigen | -2.063 | 0.872 | 1.541 | 344115038 | 136 |
| CAPG | Macrophage-capping protein | -2.063 | 0.777 | 1.541 | 352255023 | 137 |
| CCL27 | C-C motif chemokine 27 | -2.23 | 0.823 | 0.994 | 359256696 | 138 |
| TDGF1 | Teratocarcinoma-derived growth factor 1 | -2.042 | 0.809 | 1.541 | 371548086 | 139 |
| HS6ST1 | Heparan-sulfate 6-O-sulfotransferase 1 | -2.48 | 0.802 | 0.619 | 381908068 | 140 |
| PRSS3 | Trypsin-3 | -2.205 | 0.823 | 0.994 | 389691303 | 141 |
| F9.1 | Coagulation factor IXab | -2.721 | 0.794 | 0 | 407565957 | 142 |
| ADAMTS15 | A disintegrin and metalloproteinase with thrombospondin motifs 15 | -2.102 | 0.834 | 1.541 | 410591411 | 143 |
| OSM | Oncostatin-M | -2.242 | 0.844 | 0.994 | 426248646 | 144 |
| ANP32B | Acidic leucine-rich nuclear phosphoprotein 32 family member B | -2.286 | 0.818 | 0.994 | 426866823 | 145 |
| CAMK1D | Calcium/calmodulin-dependent protein kinase type 1D | -2.084 | 0.844 | 1.541 | 444783686 | 146 |
| F10 | Coagulation factor Xa | -2.4 | 0.812 | 0.619 | 447143476 | 147 |

| | | | | | | |
|---|---|---|---|---|---|---|
| C8A.C8B.<br>C8G | Complement component C8 | -2.154 | 0.849 | 0.994 | 459197271 | 148 |
| IL37 | Interleukin-37 | -2.274 | 0.828 | 0.994 | 466347158 | 149 |
| IFNL2 | Interferon lambda-2 | -2.128 | 0.831 | 1.541 | 484511388 | 150 |
| F9 | Coagulation factor IX | -2.189 | 0.845 | 0.994 | 487734706 | 151 |

**Table S5 Single Etiology validation statistics for BvV and BvVM marker sets** (A to D)**, and SOMAScan assay and RNA-Sequencing overlapping marker set (n=24) for RF and EN models** (E and F)

**(A) Single Etiology validation set (n=32) for Bacteria vs Virus (Random Forest Model).** Performance statistics summary for 5, 10, 15, 25, 50, 100, and 219 markers. The validation set (n=32) contains Bacteria (n=19) and Virus (n=13). The confusion matrix and other statistics are included in Table 1.

| Model | Bacteria vs. Virus (n=32) Random Forest | | | | | | |
|---|---|---|---|---|---|---|---|
| # of markers | All (219) | 100 | 50 | 25 | 15 | 10 | 5 |
| Sensitivity | 0.89 | 0.95 | 0.95 | 0.89 | 0.84 | 0.84 | 0.89 |
| Specificity | 1.00 | 0.92 | 1.00 | 0.92 | 0.85 | 0.85 | 0.85 |

**(B) Single Etiology validation set (n=47) for Bacteria vs Virus or Malaria (Random Forest Model).** Performance statistics summary for 5, 10, 15, 25, 50, 100, and 151 markers. The validation set (n=51) contains Bacteria (n=19), Malaria (n=15), and Virus (n=13). The confusion matrix and other statistics are included in Table 1.

| Model | Bacteria vs. Virus or Malaria (n=47) Random Forest | | | | | | |
|---|---|---|---|---|---|---|---|
| # of markers | All (151) | 100 | 50 | 25 | 15 | 10 | 5 |
| Sensitivity | 0.68 | 0.74 | 0.63 | 0.74 | 0.63 | 0.58 | 0.53 |
| Specificity | 1.00 | 0.89 | 0.93 | 0.93 | 0.86 | 0.86 | 0.64 |

**(C) Single Etiology validation set (n=32) for Bacteria vs Virus (Elastic Net Model).** Confusion matrices with all (219) and 5 markers and performance statistics summary for 5, 10,15, 25, 50, 100, and 219 markers including accuracy with 0%, 10% and 20% surrogates. The validation set (n=32) contains Bacteria (n=19) and Virus (n=13).

| **Model** | **Bacteria vs. Virus (n=32)** Elastic net | | | | | | | |
|---|---|---|---|---|---|---|---|---|
| # of markers | **all significant markers (219)** | | | | **5 markers** | | | |
| **Confusion Matrix** | | | *Actual* Bacteria | *Actual* Virus | | | *Actual* Bacteria | *Actual* Virus |
| | *Predicted* | Bacteria | 16 | 1 | *Predicted* | Bacteria | 18 | 4 |
| | *Predicted* | Virus | 3 | 12 | *Predicted* | Virus | 1 | 9 |
| **Accuracy** | 0.88 | | | | 0.84 | | | |
| **95% CI** | (0.71, 0.96) | | | | (0.67, 0.95) | | | |
| **Sensitivity** | 0.84 | | | | 0.95 | | | |
| **Specificity** | 0.92 | | | | 0.69 | | | |

| # of markers | **All (219)** | **100** | **50** | **25** | **15** | **10** | **5** |
|---|---|---|---|---|---|---|---|
| **Accuracy** | 0.88 | 0.84 | 0.88 | 0.91 | 0.91 | 0.78 | 0.84 |
| 10% surrogates | 0.94 | 0.84 | 0.88 | 0.91 | 0.91 | 0.88 | 0.84 |
| 20% surrogates | 0.84 | 0.81 | 0.88 | 0.88 | 0.91 | 0.88 | 0.81 |
| **Sensitivity** | 0.84 | 0.79 | 0.84 | 0.95 | 0.89 | 0.79 | 0.95 |
| **Specificity** | 0.92 | 0.92 | 0.92 | 0.85 | 0.92 | 0.77 | 0.69 |

**(D) Single Etiology validation set (n=47) for Bacteria vs Malaria or Virus (Elastic Net Model).** Confusion matrices with all (151) and 5 markers and performance statistics summary for 5, 10,15, 25, 50, 100, and 151 markers including accuracy with 0%, 10% and 20% surrogates. The validation set (n=47) contains Bacteria (n=19), Malaria (n=15), and Virus (n=13).

<table>
<tr><td>Model</td><td colspan="8">Bacteria vs. Virus or Malaria (n=47)<br>Elastic net</td></tr>
<tr><td># of markers</td><td colspan="4">all significant markers (151)</td><td colspan="4">5 markers</td></tr>
<tr><td rowspan="4">Confusion Matrix</td><td colspan="2"></td><td colspan="2">Actual</td><td colspan="2"></td><td colspan="2">Actual</td></tr>
<tr><td colspan="2"></td><td>Bacteria</td><td>Virus or Malaria</td><td colspan="2"></td><td>Bacteria</td><td>Virus or Malaria</td></tr>
<tr><td rowspan="2">Predicted</td><td>Bacteria</td><td>18</td><td>0</td><td rowspan="2">Predicted</td><td>Bacteria</td><td>10</td><td>3</td></tr>
<tr><td>Virus or Malaria</td><td>1</td><td>28</td><td>Virus or Malaria</td><td>9</td><td>25</td></tr>
<tr><td>Accuracy</td><td colspan="4">0.98</td><td colspan="4">0.74</td></tr>
<tr><td>95% CI</td><td colspan="4">(0.89, 1.00)</td><td colspan="4">(0.60, 0.86)</td></tr>
<tr><td>Sensitivity</td><td colspan="4">0.95</td><td colspan="4">0.53</td></tr>
<tr><td>Specificity</td><td colspan="4">1.00</td><td colspan="4">0.89</td></tr>
</table>

| # of markers | All (151) | 100 | 50 | 25 | 15 | 10 | 5 |
|---|---|---|---|---|---|---|---|
| **Accuracy** | 0.98 | 0.72 | 0.83 | 0.85 | 0.77 | 0.79 | 0.74 |
| 10% surrogates | 0.91 | 0.83 | 0.83 | 0.74 | 0.81 | 0.79 | 0.74 |
| 20% surrogates | 0.91 | 0.83 | 0.83 | 0.77 | 0.83 | 0.79 | 0.74 |
| **Sensitivity** | 0.95 | 0.58 | 0.63 | 0.74 | 0.53 | 0.58 | 0.53 |
| **Specificity** | 1.00 | 0.82 | 0.96 | 0.93 | 0.93 | 0.93 | 0.89 |

**(E) Single Etiology SOMAScan assay and RNA-Sequencing overlapping marker sets (n=24,** see Fig. 4 B) **predicted performance in the Bacteria vs Virus marker set (RF and EN models).** Confusion matrices and performance statistics summary depicted. The sample set (n=32) contains Bacteria (n=19) and Virus (n=13).

| **Model** | **Bacteria vs. Virus (n=32)** Random Forest | | | **Bacteria vs. Virus (n=32)** Elastic Net | | |
|---|---|---|---|---|---|---|
| # of markers | **RNA-Seq / SOMASCan (24 markers)** | | | **RNA-Seq / SOMASCan (24 markers)** | | |
| **Confusion Matrix** | | *Actual* | | | *Actual* | |
| | | Bacteria | Virus | | Bacteria | Virus |
| | *Predicted* Bacteria | 14 | 2 | *Predicted* Bacteria | 16 | 1 |
| | *Predicted* Virus | 5 | 11 | *Predicted* Virus | 3 | 12 |
| **Accuracy** | 0.78 | | | 0.88 | | |
| **95% CI** | (0.60, 0.91) | | | (0.71, 0.96) | | |
| **Sensitivity** | 0.74 | | | 0.84 | | |
| **Specificity** | 0.85 | | | 0.92 | | |

**(F) Single Etiology SOMAScan assay and RNA-Sequencing overlapping marker sets (n=24,** see Fig. 4 B**) predicted performance in the Bacteria vs Malaria or Virus sample set (RF and EN models).** Confusion matrices and performance statistics summary depicted. The validation set (n=47) contains Bacteria (n=19), Malaria (n=15), and Virus (n=13).

<table>
<tr><th>Model</th><th colspan="4">Bacteria vs. Virus or Malaria (n=47)<br>Random Forest</th><th colspan="4">Bacteria vs. Virus or Malaria (n=47)<br>Elastic Net</th></tr>
<tr><td># of markers</td><td colspan="4">RNA-Seq / SOMASCan (24 markers)</td><td colspan="4">RNA-Seq / SOMASCan (24 markers)</td></tr>
<tr><td rowspan="4">Confusion Matrix</td><td colspan="2" rowspan="2"></td><td colspan="2">Actual</td><td colspan="2" rowspan="2"></td><td colspan="2">Actual</td></tr>
<tr><td>Bacteria</td><td>Virus or Malaria</td><td>Bacteria</td><td>Virus or Malaria</td></tr>
<tr><td rowspan="2">Predicted</td><td>Bacteria</td><td>12</td><td>2</td><td rowspan="2">Predicted</td><td>Bacteria</td><td>14</td><td>2</td></tr>
<tr><td>Virus or Malaria</td><td>7</td><td>26</td><td>Virus or Malaria</td><td>5</td><td>26</td></tr>
<tr><td>Accuracy</td><td colspan="4">0.81</td><td colspan="4">0.85</td></tr>
<tr><td>95% CI</td><td colspan="4">(0.67, 0.91)</td><td colspan="4">(0.72, 0.94)</td></tr>
<tr><td>Sensitivity</td><td colspan="4">0.63</td><td colspan="4">0.74</td></tr>
<tr><td>Specificity</td><td colspan="4">0.93</td><td colspan="4">0.93</td></tr>
</table>

**Table S6 Mixed infection set without controls (n=23) (Random Forest)**

The mixed infection test set (n=23) contains the following samples: Bacteria & Malaria (n=9), Virus and probable Bacterial secondary co-infection (n=11), Malaria & Other (n=1), Malaria & Virus (n=2). All samples that contain the term "bacteria" are considered bacteria-positive samples.

**(A) Mixed infection test samples (n=23) for Bacteria vs Virus.** Confusion matrices with 219 and 5 markers and performance statistics summary for 5, 10, 15, 25, 50, 100, and 219 markers.

<table>
<tr><th>Model</th><th colspan="8">Bacteria vs. Virus (n=23)<br>Random Forest</th></tr>
<tr><td># of markers</td><td colspan="4">all significant markers (219)</td><td colspan="4">5 markers</td></tr>
<tr><td rowspan="4">Confusion Matrix</td><td colspan="2" rowspan="2"></td><td colspan="2">Actual</td><td colspan="2" rowspan="2"></td><td colspan="2">Actual</td></tr>
<tr><td>Bacteria</td><td>no Bacteria</td><td>Bacteria</td><td>no Bacteria</td></tr>
<tr><td rowspan="2">Predicted</td><td>Bacteria</td><td>19</td><td>3</td><td rowspan="2">Predicted</td><td>Bacteria</td><td>16</td><td>2</td></tr>
<tr><td>no Bacteria</td><td>1</td><td>0</td><td>no Bacteria</td><td>4</td><td>1</td></tr>
<tr><td>Accuracy</td><td colspan="4">0.83</td><td colspan="4">0.74</td></tr>
<tr><td>95% CI</td><td colspan="4">(0.61, 0.95)</td><td colspan="4">(0.52, 0.90)</td></tr>
<tr><td>Sensitivity</td><td colspan="4">0.95</td><td colspan="4">0.80</td></tr>
<tr><td>Specificity</td><td colspan="4">0</td><td colspan="4">0.33</td></tr>
</table>

| # of markers | **All (219)** | **100** | **50** | **25** | **15** | **10** | **5** |
|---|---|---|---|---|---|---|---|
| **Accuracy** | 0.83 | 0.83 | 0.83 | 0.83 | 0.74 | 0.74 | 0.74 |
| **Sensitivity** | 0.95 | 0.90 | 0.95 | 0.90 | 0.85 | 0.85 | 0.80 |
| **Specificity** | 0 | 0.33 | 0 | 0.33 | 0 | 0 | 0.33 |

**(B) Mixed infection test samples (n=23) for Bacteria vs Malaria or Virus.** Confusion matrices with 151 and 5 markers and performance statistics summary for 5, 10,15, 25, 50, 100, and 151 markers.

| Model | Bacteria vs. Virus or Malaria (n=23)<br>Random Forest | | | | | | | |
|---|---|---|---|---|---|---|---|---|
| # of markers | **all significant markers (151)** | | | | **5 markers** | | | |
| **Confusion Matrix** | | | *Actual* | | | | *Actual* | |
| | | | Bacteria | Virus or Malaria | | | Bacteria | Virus or Malaria |
| | *Predicted* | Bacteria | 13 | 2 | *Predicted* | Bacteria | 17 | 3 |
| | *Predicted* | Virus or Malaria | 7 | 1 | *Predicted* | Virus or Malaria | 3 | 0 |
| **Accuracy** | 0.61 | | | | 0.74 | | | |
| **95% CI** | (0.39, 0.80) | | | | (0.52, 0.90) | | | |
| **Sensitivity** | 0.65 | | | | 0.85 | | | |
| **Specificity** | 0.33 | | | | 0 | | | |

| # of markers | **All (151)** | **100** | **50** | **25** | **15** | **10** | **5** |
|---|---|---|---|---|---|---|---|
| **Accuracy** | 0.61 | 0.78 | 0.74 | 0.83 | 0.83 | 0.87 | 0.74 |
| **Sensitivity** | 0.65 | 0.80 | 0.80 | 0.90 | 0.95 | 0.95 | 0.85 |
| **Specificity** | 0.33 | 0.67 | 0.33 | 0.33 | 0 | 0.33 | 0 |

**Table S7 ALgorithm for MANagement of Childhood illnesses (ALMANACH).**

**(A) ALMANACH prediction in single etiology specimens from the test set (n=27) including bacterial (n=14) and viral (n=13) samples.** Confusion matrices based on ALMANACH metrics for pneumonia requiring antibiotic therapy, including chest indrawing alone, measured respiratory rate over 50 breaths/minute alone, or the presence of either indrawing or elevated respiratory rate. See accompanying text for details.

| **Model** | **ALMANACH clinical assessment** (n=27) | | | | | | | | | | | |
|---|---|---|---|---|---|---|---|---|---|---|---|---|
| method | **respiratory rate > 50/min** OR **chest indrawing** | | | | **respiratory rate > 50/min** | | | | **chest indrawing** | | | |
| **Confusion Matrix** | | | *Actual* | | | | *Actual* | | | | *Actual* | |
| | | | Pneumonia | non Pneumonia | | | Pneumonia | non Pneumonia | | | Pneumonia | non Pneumonia |
| | **ALMANACH** *Predicted* | Pneumonia | 13 | 10 | **ALMANACH** *Predicted* | Pneumonia | 10 | 8 | **ALMANACH** *Predicted* | Pneumonia | 12 | 7 |
| | | non Pneumonia | 1 | 3 | | non Pneumonia | 4 | 5 | | non Pneumonia | 2 | 6 |
| **Accuracy** | 0.59 | | | | 0.56 | | | | 0.67 | | | |
| **95% CI** | (0.39, 0.78) | | | | (0.35, 0.75) | | | | (0.46, 0.83) | | | |
| **NIR** | 0.52 | | | | 0.52 | | | | 0.52 | | | |
| **p-value** | 0.28 | | | | 0.43 | | | | 0.088 | | | |
| **Sensitivity** | 0.93 | | | | 0.71 | | | | 0.86 | | | |
| **Specificity** | 0.23 | | | | 0.38 | | | | 0.46 | | | |

**(B) ALMANACH clinical assessment with single etiology specimens from the test set (n=40) with bacterial (n=14), viral (n=13), and malarial (n=13) samples.** Confusion matrices based on ALMANACH metrics for pneumonia requiring antibiotic therapy, including chest indrawing alone, measured respiratory rate over 50 breaths/minute alone, or the presence of either indrawing or elevated respiratory rate. See accompanying text for details.

<table>
<tr><th>Model</th><th colspan="9">ALMANACH clinical assessment<br>(n=40)</th></tr>
<tr><td>method</td><td colspan="3">respiratory rate > 50/min<br>OR chest indrawing</td><td colspan="3">respiratory rate > 50/min</td><td colspan="3">chest indrawing</td></tr>
<tr><td rowspan="4">Confusion Matrix</td><td rowspan="2"></td><td colspan="2">Actual</td><td rowspan="2"></td><td colspan="2">Actual</td><td rowspan="2"></td><td colspan="2">Actual</td></tr>
<tr><td>Pneumonia</td><td>non Pneumonia</td><td>Pneumonia</td><td>non Pneumonia</td><td>Pneumonia</td><td>non Pneumonia</td></tr>
<tr><td>ALMANACH Predicted Pneumonia</td><td>13</td><td>17</td><td>ALMANACH Predicted Pneumonia</td><td>10</td><td>14</td><td>ALMANACH Predicted Pneumonia</td><td>12</td><td>9</td></tr>
<tr><td>ALMANACH Predicted non Pneumonia</td><td>1</td><td>9</td><td>ALMANACH Predicted non Pneumonia</td><td>4</td><td>12</td><td>ALMANACH Predicted non Pneumonia</td><td>2</td><td>19</td></tr>
<tr><td>Accuracy</td><td colspan="3">0.55</td><td colspan="3">0.55</td><td colspan="3">0.74</td></tr>
<tr><td>95% CI</td><td colspan="3">(0.38, 0.71)</td><td colspan="3">(0.38, 0.71)</td><td colspan="3">(0.58, 0.86)</td></tr>
<tr><td>NIR</td><td colspan="3">0.65</td><td colspan="3">0.65</td><td colspan="3">0.67</td></tr>
<tr><td>p-value</td><td colspan="3">0.93</td><td colspan="3">0.93</td><td colspan="3">0.21</td></tr>
<tr><td>Sensitivity</td><td colspan="3">0.93</td><td colspan="3">0.71</td><td colspan="3">0.86</td></tr>
<tr><td>Specificity</td><td colspan="3">0.35</td><td colspan="3">0.46</td><td colspan="3">0.68</td></tr>
</table>

8. Kim JW, Botvinnik OB, Abudayyeh O, et al. Characterizing genomic alterations in cancer by

complementary functional associations. Nat Biotechnol 2016;34(5):539–46.

18. Craig A, Mai J, Cai S, Jeyaseelan S. Neutrophil recruitment to the lungs during bacterial pneumonia.

Infect Immun 2009;77(2):568–75.

33. Tibshirani R, Seo MJ, Chu G, Narasimhan B, Li J, Tibshirani MR. Package “samr.” 2018;Available from: ftp://64.50.236.52/.1/cran/web/packages/samr/samr.pdf
